# Nature’s Antivenom: Combinations of conserved rattlesnake serum metalloproteinase inhibitors block the lethal action of viper venoms

**DOI:** 10.64898/2026.04.13.718210

**Authors:** Sean B. Carroll, Fiona P. Ukken, Yetunde A. Ayinuola, Luis Escalona, Montamas Suntravat, Elda E. Sanchez

**Author notes:** Corresponding author: Sean B Carroll. These authors contributed equally to this work (Sean B. Carroll, Fiona P. Ukken). Author Contributions: S.B.C. conceptualized the study, oversaw collaboration, wrote the initial draft; F.P.U. performed experiments, created presentation, reviewed and edited the manuscript; Y.A.A. performed experiments; L.E. analyzed animal experimental data; M.S. performed experiments and statistical analyses; E.E.S led and performed animal studies, reviewed and edited the manuscript. Competing Interest Statement: S.B.C. and F.P.U are co-inventors on patents pending from related work, of which the University of Maryland is the assignee (U.S. Patent App. No. 63/431,147 and 63/994,480). Classification: Biological Sciences: Medical Sciences.

## Abstract

Snakebite maims or kills several hundred thousand people each year. For more than a century, treatment has relied on antivenoms derived from animals immunized with whole venoms, but their efficacy, safety, and availability are highly variable, and it is often not well understood which specific venom components must be inhibited to prevent mortality and major morbidities. New therapeutic approaches are needed. Here, we take an evolutionary approach to antivenom design inspired by the longstanding observation that vipers have evolved serum-borne toxin inhibitors that confer resistance to their own venoms. We have investigated the abilities of a family of four rattlesnake metalloproteinase (MP) inhibitors derived from the ancestral serum glycoprotein Fetuin-A (FETUAs) to neutralize the enzymatic, hemorrhagic, and lethal activities of viper venoms. We find that while certain individual FETUA proteins are able to inhibit enzymatic or hemorrhagic activity, they are unable or only partially able to inhibit venom lethality. However, we show that specific combinations of FETUA proteins complement one another’s activities and are sufficient to fully neutralize rattlesnake venom lethality with approximately 10 times greater potency than commercial antivenom. Moreover, we demonstrate that FETUA proteins are well-conserved among viper subfamilies and that rattlesnake FETUAs are able to inhibit the MPs and neutralize the lethality of several evolutionarily distant pit viper or true viper venoms. Our results highlight the critical importance of inhibiting MPs in hemorrhagic venoms and the potential general utility of combinations of naturally evolved, recombinant MP inhibitors in the treatment of viper snakebite.

**Significance Statement:** Snakebite remains an undertreated tropical disease that affects over one million people annually, and new therapeutic approaches are needed. Here, we develop a novel approach to viper antivenoms inspired by the observations that these snakes are typically resistant to the hemorrhagic and lethal effects of their own venoms and that this resistance can be transferred to experimental animals through snake sera. We show that a small set of serum metalloproteinase inhibitors from the Western Diamondback rattlesnake are potent inhibitors of the lethality of its own venom, and able to block the lethality of other rattlesnakes and evolutionarily distant vipers. Recombinant natural antitoxins offer a promising approach for formulating more potent viper antivenoms.

## Introduction

Snakebite affects at least 1.7 million people annually, causing approximately 100,000 deaths and 400,000 disabilities(1), an impact that can exceed that of tropical diseases such as malaria, dengue, or trypanosomiasis in certain areas(2) . The current standard of therapy involves the administration of antivenoms composed of polyclonal immunoglobulin preparations from large animals that have been hyperimmunized with whole venoms of snake species found in a given region. However, in many parts of the world snakebite treatment is inadequate due to the lack of access to effective and safe antivenoms (3).

Some of the root causes of the low quality of antivenoms are due to limitations inherent to the current method of antivenom production, which has not changed much since its introduction in the late 19^th^ century (1, 4). Two critical parameters of antivenom efficacy are their potency – the dose required to neutralize a given venom’s toxic effects, and their polyvalency - the ability to effectively neutralize the spectrum of venomous species found within a target geographic area. Optimization of these two parameters is made especially challenging by the biochemical complexity and diversity of snake venoms.

The most dangerous and medically important snakes belong to two major families: the Elapidae (elapids, including coral snakes, cobras, kraits, mambas, tiger snakes and brown snakes) and the Viperidae (true vipers and pit vipers). An individual venom may contain as many as 50-100 toxin proteins belonging to a dozen or more protein families (5, 6), and any two species’ venoms can differ considerably in composition with the scope of such differences generally increasing with their evolutionary and ecological divergence (7–9). The selection of immunizing venoms to increase polyvalency may sacrifice potency, and vice versa, yielding antivenoms with relatively low activity that need to be administered in larger doses. Moreover, antivenom potency is limited by the method used for purification: almost all antivenoms are total immunoglobulin preparations in which only a small percentage of the antivenom protein consists of antibodies specific to venom toxins (10). The large doses required for low-purity, low-potency, animal-derived antivenoms in turn increases the risk of side effects such as anaphylaxis and serum sickness(11) as well as the costs of treatment and the pressures on limited antivenom supplies.

The continuing high incidence of snakebite mortality and morbidity and persistent problems with antivenom quality and availability led the World Health Organization in 2017 to declare snakebite a Neglected Tropical Disease and to call for a series of measures to address the burdens of snakebite across the globe, including innovations to improve the efficacy, consistency, and availability of antivenoms (3).

A few new strategies have recently emerged to potentially address the shortcomings of current antivenoms including: i) small molecule inhibitors that target major classes of venoms toxins such as the orally available metalloproteinase inhibitor marimastat and phospholipase A2 inhibitor varespladib that may be delivered to victims in the field before reaching a hospital (12–17); ii) monoclonal antibodies and nanobodies raised against individual venom toxins and selected for their neutralizing power and species cross-reactivity and which have shown particular promise against neurotoxic elapid venoms including cobras, kraits, and mambas in animal studies(18–21) and iii) de novo designed protein antitoxins developed using molecular modeling and deep learning methods (22).

We are exploring an alternative strategy for antivenom design inspired by the long-standing observations that some snakes, particularly vipers and pit vipers, exhibit resistance to their own and other species’ venoms (23–25) and that their sera can protect experimental animals from lethal effects (26–29). Several serum-borne inhibitors have been isolated and characterized largely from pit vipers and target toxins that are members of the group II phospholipase A2 (PLA_2_) (30) or metalloproteinase (MP) protein families.

The MP protein family is primarily responsible for hemorrhage, tissue damage, and coagulopathy (31) and is typically the most abundant toxin family in pit viper venoms (∼50% of venom protein by weight in *C. atrox* (32)). It is also the largest toxin family (with thirty members in *C. atrox*; (33)) and the most structurally diverse, consisting of three main classes of enzymes constructed of different combinations of class-defining protein domains: P-I class MPs (MPO) possess only the metalloproteinase domain; P-II class MPs (MAD) possess the metalloproteinase and disintegrin domains; and P-III class MPs (MDC) possess the metalloproteinase, disintegrin, and cysteine-rich domains (reviewed in (34). The most-studied serum metalloproteinase inhibitor has been isolated from the sera of three pit viper species, *Protobothrops flavoviridis* (Habu Serum Factor, HSF (35)), *Gloydius blomhoffi* (Mamushi serum factor, MSF (36)), and *Bothrops jararaca* (Bj46a; (37)) on the basis of its ability to inhibit venom hemorrhagic activity and shown to be a cystatin-type MP inhibitor related to the serum protein Fetuin-A (we refer to this protein as FETUA-2 and the family as FETUA proteins (38).

It is not known, however, how many different specific toxin inhibitors exist in pit viper sera, nor which inhibitors are necessary to inhibit venom lethality as the lethality-neutralizing activity of whole sera against hemorrhagic venoms has not been reproduced with isolated serum proteins. We recently discovered that the FETUA protein family in rattlesnakes and other pit vipers is larger than previously known, identified five members in the rattlesnake genus *Crotalus*, and showed that one *C. atrox* protein (FETUA-3) inhibited three distinct classes of venom MPs in vitro (38). However, the roles of the different FETUA proteins, whether they act as general MP inhibitors or inhibitors of subsets or of specific MPs, are not known. Here, we have investigated the abilities of *C. atrox* FETUA proteins to inhibit the activities of rattlesnake venoms in vivo. We find that individual FETUA proteins are either ineffective or only partially effective at inhibiting venom lethality, whereas specific combinations of FETUA proteins are sufficient to fully neutralize venom lethality with approximately ten times the potency of affinity-purified commercial antivenom. We also demonstrate that critical FETUA proteins are deeply conserved among vipers and can inhibit MPs and neutralize the lethality of venoms from evolutionarily distant species. We suggest that combinations of recombinant MP inhibitors offer a new, potentially general approach to formulating potent, polyvalent viper antivenoms.

## Results

### *Crotalus atrox* serum is a potent inhibitor of venom lethality

The ability of rattlesnake sera to neutralize their own venoms was reported many decades ago(26, 29). To obtain a measure of the potency of rattlesnake sera relative to antivenom, we compared the specific activity of whole *C. atrox* serum with that of the commercial North American pit viper antivenom CroFab, the only antivenom comprised exclusively of affinity-purified, venom-toxin binding antibodies that is obtained from animals immunized with four pit viper venoms, including *C. atrox* (39). In a standard antivenom lethality neutralization assay in which antivenom or serum is pre-incubated with 3 x LD_50_ of *C. atrox* venom (LD_50_ 4.1mg/kg), and injected intraperitoneally into mice, we determined a CroFab antivenom median effective dose (ED_50_) of 52.6 ± 41.5 mg/kg and a *C. atrox* serum ED_50_ of 17.1 ± 4.7 mg/kg. The greater potency of whole, unfractionated *C. atrox* serum relative to the affinity-purified antivenom suggests that the rattlesnake serum contains potent inhibitors of venom toxins that are able, individually or collectively, to counteract the lethal action of the venom. Since MPs are responsible for the hemorrhage and tissue destruction that are hallmarks of *C. atrox* envenomation (31), and the most abundant toxin family in the venom (32), we focused on potential inhibitors of the MP family.

### FETUA proteins differ in their ability to inhibit *C. atrox* venom hemorrhagic activity

We previously identified five FETUA proteins in *C. atrox* that are shared among *Crotalus* species (38). FETUA-1 is the ancestral-type Fetuin-A shared with all snakes and other vertebrates whereas FETUA-2, 3, 4, and 5 all evolved within the lineage leading to rattlesnakes. FETUA-1 is a two-chain molecule due to post-translational cleavage whereas FETUA-2,3,4, and 5 are all single-chain molecules due to the loss of the ancestral cleavage site (Fig. 1A,B). Each FETUA paralog has a signature N-terminal sequence that distinguishes it from the other paralogs (Fig.1A). FETUA-1, FETUA-2, and FETUA-4 do not inhibit whole *C. atrox* venom or purified MP collagenase activity. FETUA-5 only partially inhibits whole venom collagenase activity, but FETUA-3 effectively inhibits whole venom collagenase activity as well as the activities of three of the most abundant MPs in *C. atrox* venom (38). Therefore, our initial in vivo experiments focused on FETUA-3 inhibition of venom hemorrhagic and lethal activities.

**Figure 1.**
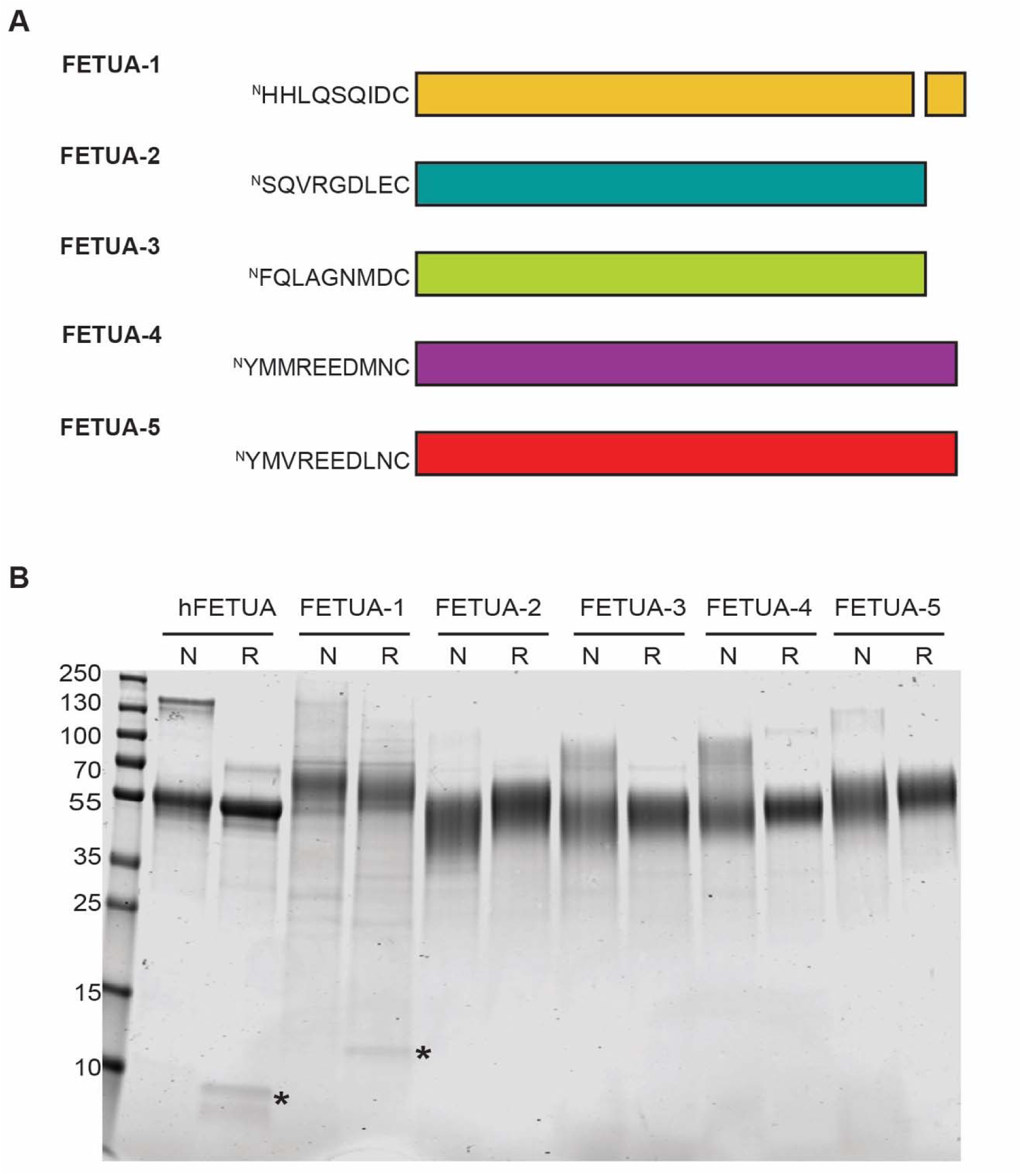
The novel *Crotalu*s FETUA proteins are single-chain molecules distinguished by their N-termini. A) Schematic of the *C. atrox* FETUA proteins. The FETUA-1 protein is the ortholog of the vertebrate ancestral Fetuin-A protein. The protein is cleaved post-translationally near the C-terminus and small light chain and large heavy chain are held together by a disulfide bond (denoted by short black line connecting two part of the molecule. FETUA2-5 are single-chain molecules, and each ortholog is distinguished by the sequences of the first nine or ten N-terminal amino acids which tend to be highly conserved among orthologs (Ukken et al. 2022). B) SDS-PAGE of native human Fetuin-A and recombinant *C. atrox* FETUA proteins under non-reducing (N) and reducing (R) conditions. Note the small peptide that appears upon reduction of human Fetuin-A and *C. atrox* FETUA-1 (asterisks); this is the short C-terminal light chain peptide that is generated post-translationally by an endopeptidase and is linked to the heavy chain by a disulfide bond. The FETUA-2-5 proteins lack this cleavage site and are single chain molecules. The FETUA-2-5 proteins exhibit the atypical behavior of migrating more slowly after reduction, perhaps due to becoming less compact after the breakage of six intramolecular disulfide bonds. The apparent molecular weights of the FETUA glycoproteins are approximately 55Kda; the mature proteins are 305-325 amino acids in length (deduced molecular weights ∼34-36 Kda). The protein amounts loaded were hFETUA and FETUA-1 (8µg) and FETUA-2-5 (5µg). The recombinant FETUA proteins used in this study are estimated to be >95% pure.

We found, however, that 10µg of recombinant FETUA-3 protein, an amount far exceeding that necessary to fully inhibit the collagenase activity of the quantity of whole venom tested, did not inhibit hemorrhagic activity when pre-incubated with a minimum hemorrhagic dose (MHD; 1.5µg) of *C. atrox* venom and injected subcutaneously into mice in a standard anti-hemorrhage assay (40)(Fig. 2A,Table 1, Fig. S1A). This result suggests that FETUA-3 does not inhibit MPs that contribute most significantly to hemorrhage in vivo.

**Figure 2.**
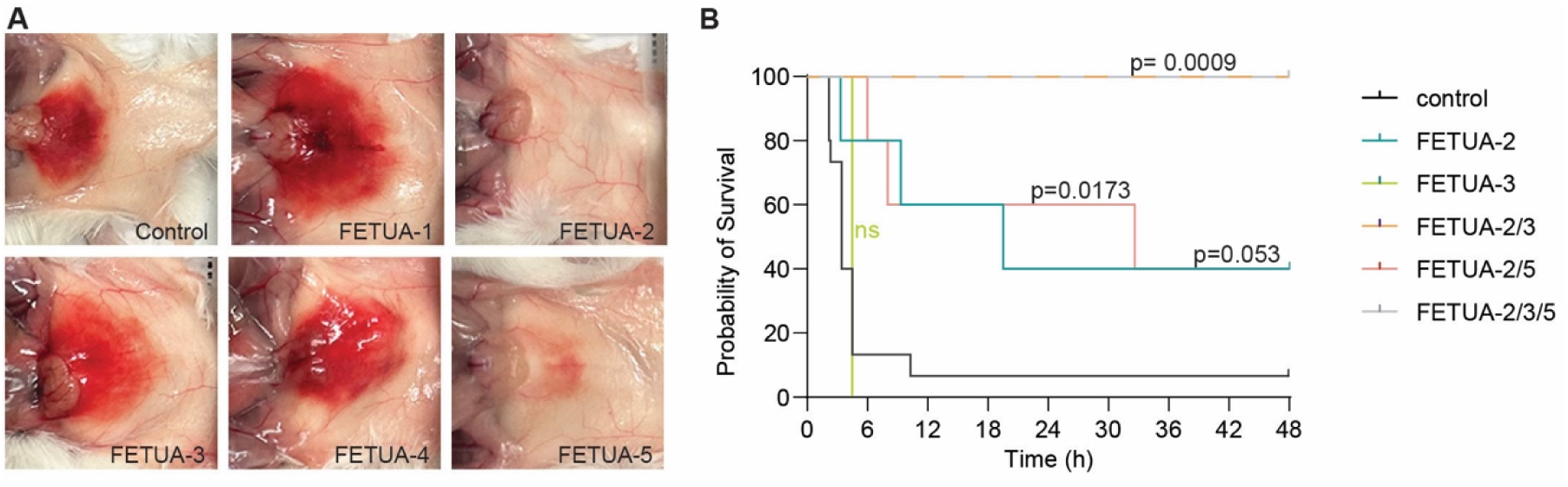
Inhibition of *C. atrox* venom hemorrhagic and lethal activities by FETUA proteins. **A.** *C. atrox* venom hemorrhagic activity is completely inhibited by FETUA-2 and strongly inhibited by FETUA-5. FETUA-1, FETUA-3, and FETUA-4 enhance hemorrhage at the high protein concentration tested. Six groups of mice (n=4 per group) were injected subcutaneously with 1.5 µg of *C. atrox* venom that was pre-incubated with either saline (control) or 10µg of individual FETUA proteins indicated. The animals were sacrificed after 24 hr, the hemorrhage was photographed and hemorrhagic units (HaU) quantified by their area and intensity. Each photograph is representative of the average HaU of each group of four mice. B. Kaplan-Meier survival plots of mice that were injected i.p. with 3 x LD50 of *C. atrox* venom that was preincubated with either saline (control), or 100µg of each individual or combination of FETUA proteins indicated, and monitored for 48 hr. Control mice died within 2-12 hrs. FETUA-2 alone partially protected mice while FETUA-3 alone provided no protection. FETUA-2 combined with FETUA-5 provided the same protection from lethality as FETUA-2 alone. However, the combination of FETUA-2 and FETUA-3 or of FETUA-2/FETUA-3/FETUA-5 provided complete protection against the lethal effect of the venom.

**Table 1.** Inhibition of *C. atrox* Venom Hemorrhagic Activity by FETUA Proteins*.

| FETUA Protein | Inhibition of Hemorrhagic Activity (%) |
| --- | --- |
| FETUA-1 | -101±30<sup>#</sup> |
| FETUA-2 | 100±0 |
| FETUA-3 | -66± 8<sup>#</sup> |
| FETUA-4 | -95±21<sup>#</sup> |
| FETUA-5 | 92±3 |

\*Four mice each were injected with 1.5 μg of *C. atrox* venom, a dose pre-determined to generate a ~10mm hemorrhagic spot (defined as the minimum hemorrhagic dose MHD), pre-mixed with 10 μg of each FETUA protein, administered subcutaneously and observed for 24hr. Hemorrhagic activity was quantified based on the area and intensity of the hemorrhagic spot (see Materials and Methods) and percent inhibition was calculated relative to the venom control (117.6 ± 37.8 HaU).
<sup>#</sup>Hemorrhagic activity was increased relative to the venom control so the value is negative.

Furthermore, 100µg of FETUA-3 protein did not prevent or delay mortality when pre-incubated with three times the LD_50_ of *C. atrox* venom and injected intraperitoneally into mice in a standard antivenom assay (40)(Fig. 2B, Table 2). The lack of even a delay in venom action by FETUA-3 suggests that some critical toxin(s) that contribute to lethality, which could be MPs and/or other toxin types, are not impeded by FETUA-3.

**Table 2.** Inhibition of *C. atrox* Venom Lethal Activity by FETUA Proteins.

| FETUA Protein | Lethal Activity #<br>(Live/Dead Mice)(P value) |
| --- | --- |
| Venom Control Only | 1/9 |
| FETUA-1 | n.t |
| FETUA-2 | 2<sup>†</sup>/3 (P =0.053) |
| FETUA-3 | 0/5 |
| FETUA-4 | n.t. |
| FETUA-5 | n.t |
| FETUA-2, FETUA-5 | 2<sup>†</sup>/3 (P = 0.017) |
| FETUA-2, FETUA-3 | 5/0 (P < 0.001) |
| FETUA-2, FETUA-3, FETUA-5 | 5/0 (P < 0.001) |

#Mice were injected intraperitoneally with 3xLD50 of *C. atrox* venom that had been pre-incubated with saline, or with 100µg of each FETUA protein. Animals were observed for 48-hours. The P value was derived from a log-rank test comparing the treatment groups with the control group.

Since FETUA-3 alone is not sufficient to inhibit either *C. atrox* venom hemorrhagic or lethal activity, we examined the ability of other FETUA proteins to neutralize these activities. To detect whether any FETUAs are capable of inhibiting hemorrhage to any degree, each FETUA protein was first tested at a large excess of protein relative to whole venom (10µg:1.5µg). We found that FETUA-2 completely inhibited hemorrhage and FETUA-5 strongly inhibited hemorrhage (Fig. 2A, Table 1; Fig. S1A).

We also observed that FETUA-1 and FETUA-4 exhibited no inhibition of hemorrhagic activity rather, like FETUA-3, they enhanced hemorrhage (Fig. 2A; Table 1; Fig. S1A). We note that we have previously observed enhancement of purified MP activity in vitro by selected FETUAs (38)), but we do not understand the mechanism of enhancement.

### The complementary action of two FETUA proteins neutralizes *C. atrox* venom lethality

The strong anti-hemorrhagic activities of FETUA-2 and FETUA-5 indicate that these proteins act on the MPs primarily responsible for causing hemorrhage. Since hemorrhage is major manifestation of *C. atrox* and most pit viper envenomations, we explored the potential abilities of these proteins to neutralize the lethal effect of *C. atrox* venom when administered in amounts sufficient to neutralize hemorrhage. We found that FETUA-2 protein alone provided some partial protection against lethality (three out of five mice died while two survivors exhibited symptoms of envenomation; Fig. 2B; P = 0.053; Table 2). This result indicates that blocking hemorrhagic MPs alone reduces but is not sufficient to fully prevent C*. atrox* venom lethality. It raises the possibility that additional MPs not inhibited by FETUA-2, or other toxin types, must also be neutralized.

To explore whether the addition of FETUA-5 might further enhance protection against lethality, we combined FETUA-5 and FETUA-2 and tested their activity in a lethality assay. We again observed only partial protection, similar to treatment with FETUA-2 alone (Fig. 2B, P = 0.017; Table 2).

Since the inclusion of FETUA-5 had no discernible additive effect on venom neutralization, we next considered whether FETUA-3, despite having no effect on hemorrhagic activity, might contribute to overall venom neutralization. Indeed, when we combined FETUA-3 and FETUA-2, we observed complete neutralization of venom lethality with all mice surviving and showing no overt symptoms of envenomation (Fig. 2B; P < 0.001 relative to control, P = 0.049 relative to FETUA-2/5 group; Table 2). We also observed complete protection with the triple combination of FETUA-2, FETUA-3, and FETUA-5 (Fig. 2B, P < 0.001; Table 2).

These results indicate that FETUA-3, which strongly inhibits whole venom MP collagenase activity but not hemorrhage, and FETUA-2, which does not inhibit whole venom collagenase activity but strongly inhibits hemorrhage, act in a complementary manner to block a spectrum of MP activities in *C. atrox* venom. Most importantly, these results also demonstrate that blocking MP activity alone is sufficient to prevent the lethal effects of *C. atrox* venom in mice, an observation not previously reported for *C. atrox* or any pit viper venom. In light of the FETUAs unexpected and unprecedented neutralizing activity, we tested whether the proteins might neutralizing any other major venom toxins but we observed no inhibitory activity against either venom serine proteases or venom phospholipase A2 (Fig. S2A,B).

### A distinct combination of FETUA proteins inhibits *Crotalus adamanteus* venom hemorrhagic and lethal activities

In light of this unexpected observation, we sought to determine whether FETUA proteins might also act in a combined and complementary manner on other rattlesnake species venoms. The gene complex encoding MP toxins has diversified rapidly among rattlesnakes such that different species encode different numbers of MP genes (5-30 genes;(33, 41–44)) and express different combinations and quantities of individual MP proteins in their venoms(33, 44). We focused on the Eastern Diamondback rattlesnake (*C. adamanteus*) because it diverged from its last common ancestor shared with *C. atrox* ∼5.5-7 mya (45, 46), produces a strongly hemorrhagic venom (47), expresses a distinct subset of its twenty-three MP genes relative to *C. atrox* (44), and this species’ venom has long been used (with *C. atrox*) as one of the primary venoms to produce and standardize North American pit viper antivenoms (39).

*C. adamanteus* FETUA protein sequences are highly similar to their *C. atrox* orthologs, such that we inferred that they are likely to be functionally equivalent (38). Therefore, we tested the *C. atrox* versions of the five FETUA proteins for their ability to inhibit *C. adamanteus* venom-induced hemorrhage. To determine whether any individual FETUAs were capable of inhibiting hemorrhage, each protein was tested at a large excess of FETUA protein relative to whole venom (10µg:1.5µg). We found again that FETUA-1, FETUA-3, and FETUA-4 did not inhibit but rather enhanced hemorrhage, however both FETUA-2 and FETUA-5 provided partial inhibition of hemorrhagic activity. In contrast to their respective action on *C. atrox* venom, FETUA-5 exhibited the stronger inhibitory activity on *C. adamanteus* venom (Fig. 3A; Table 3; Fig. S1B).

**Figure 3.**
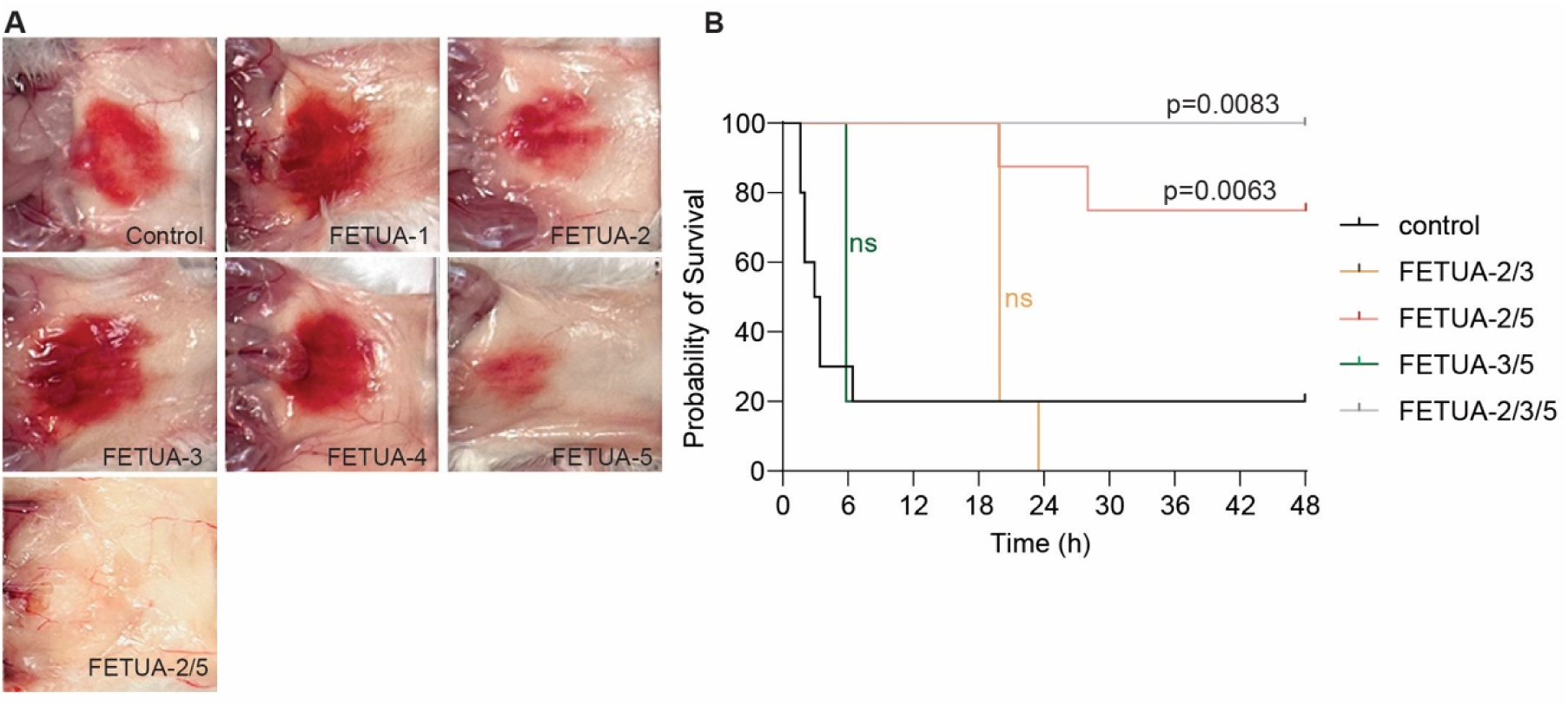
Inhibition of *C. adamanteus* venom hemorrhagic and lethal activities by FETUA proteins. **A**. *C. adamanteus* venom hemorrhagic activity is partially inhibited by FETUA-2 and FETUA-5, and completely inhibited by the combination of FETUA-2 and FETUA-5. FETUA-1, FETUA-3, and FETUA-4 enhance hemorrhage at the high protein concentration tested. Seven groups of mice (n=4 per group) were injected subcutaneously with 1.5 µg of *C. adamanteus* venom that was pre-incubated with saline (control) or 10µg of individual FETUA proteins indicated. The animals were sacrificed after 24 hr, the hemorrhage was photographed and hemorrhagic units (HaU) quantified by their area and intensity. Each photograph is representative of the average HaU of each group of four mice. B. Kaplan-Meier survival plots of mice that were injected i.p. with 3 x LD50 of *C.adamanteus* venom that was preincubated with either saline (control), or 100µg of each FETUA protein in the combinations indicated, and monitored for 48 hr. Control mice died within approximately 6 hr, we interpret survivors as having been mis-injected. The combination of FETUA-2 and FETUA-3 provided no protection, nor did the combination of FETUA-3 and FETUA-5 (same survival as control). The combination of FETUA-2 and FETUA-5 provided substantial, but incomplete protection, while the combination of FETUA-2/3/5 provided complete protection against venom lethality.

**Table 3.** Inhibition of *C. adamanteus* Venom Hemorrhagic Activity by FETUA Proteins*.

| FETUA Protein | Inhibition of Hemorrhagic Activity (%) |
| --- | --- |
| FETUA-1 | -119±31<sup>#</sup> |
| FETUA-2 | 29±9 |
| FETUA-3 | -161±22<sup>#</sup> |
| FETUA-4 | -51±7<sup>#</sup> |
| FETUA-5 | 61±4 |
| FETUA- 2, FETUA-5 | 100±0 |

\*Four mice each were injected with 1.5 μg of *C. adamanteus* venom, a dose pre-determined to generate a ~10mm hemorrhagic spot (defined as the minimum hemorrhagic dose MHD), pre-mixed with 10 μg of each FETUA protein, administered subcutaneously and observed for 24hrs. Hemorrhagic activity was quantified based on the area and intensity of the hemorrhagic spot (see Materials and Methods) and percent inhibition was calculated relative to the venom control (97.9 ± 10.4 HaU).
<sup>#</sup>Hemorrhagic activity was increased relative to the venom control so the value is negative.

The incomplete inhibition of *C. adamanteus* hemorrhagic activity by both FETUA-2 and by FETUA-5 could be due to a variety of factors including weaker interactions between the FETUAs and hemorrhage-causing *C. adamanteus* MPs, redundant activity of the two FETUAs on the same subset of MPs, or non-overlapping activities of each FETUA where each inhibits a non-identical subset of hemorrhagic MP(s) such that treatment with one protein leaves some hemorrhagic MP(s) uninhibited. To attempt to resolve among these possibilities, we tested the effect of combining FETUA-2 and FETUA-5 in the hemorrhage assay and observed complete inhibition of hemorrhage (Fig. 3A; Table 3; Fig. S1B). This result is consistent with the two proteins acting in a complementary fashion to inhibit different hemorrhage-causing MPs.

We then tested whether FETUA proteins were able to inhibit the lethal effects of *C. adamanteus* venom. Because of their individual and combined complementary effects in inhibiting hemorrhage, we focused on the FETUA-2 and FETUA-5 proteins and tested various combinations of these proteins with and without FETUA-3 for their ability to neutralize venom lethality. In striking contrast to their ability to neutralize *C. atrox* venom, we observed no neutralization of *C. adamanteus* venom by the combination of FETUA-2 and FETUA-3 (Fig. 3B; P = 0.27; Table 4). The combination of FETUA-3 and FETUA-5 together also did not neutralize C*. adamanteus* venom (Fig. 3B; Table 4). The combination of FETUA-2 and FETUA-5 together provided substantial but incomplete protection against lethality (Fig. 3B, P = 0.0063; Table 4). However, the combination of FETUA-3 with FETUA-2 and FETUA-5 provided complete protection (Fig. 3B, P = 0.0083; Table 4).

**Table 4.** Inhibition of *C. adamanteus* Venom Lethal Activity by Combinations of FETUA Proteins^#^.

| FETUA Proteins | Lethal Activity<br>(Live/Dead mice) |
| --- | --- |
| Venom Control Only | 2/8 |
| FETUA-2, FETUA-3 | 0/5 |
| FETUA-3, FETUA-5 | 1/4 |
| FETUA-2, FETUA-5 | 6/2 (P =0.0063) |
| FETUA-2, FETUA-3, FETUA-5 | 5/0 (P =0.0083) |

<sup>#</sup>Mice were injected intraperitoneally with 3xLD<sub>50</sub> of *C. adamanteus* venom that had been pre-incubated with saline, or a mixture containing 200µg of each of the FETUA proteins. Animals were observed for 48hrs. The P value was derived from a log-rank test comparing the treatment groups with the control group.

These results with *C. adamanteus* venom provide additional support that blocking MP activity is sufficient to neutralize the lethal activity of hemorrhagic rattlesnake venoms. They also reveal that different FETUA protein combinations are required to neutralize different species’ venoms, with FETUA-2/FETUA-3 sufficient to effectively neutralize *C. atrox* but not *C. adamanteus* venom, and FETUA-2/FETUA-3/ FETUA-5 required to fully neutralize *C. adamanteus* venom.

To compare the relative potency of the FETUA-2/FETUA-3/FETUA-5 protein combination with those previously determined above for *C. atrox* sera and commercial CroFab antivenom, we pre-incubated varying amounts of the FETUA combination with three times the LD_50_ of *C. atrox* venom and injected the mixtures intraperitoneally into mice. We obtained an ED_50_ for the FETUA combination of just 5.6 ­± 1.9 mg/kg, which is approximately three times the potency of *C. atrox* sera (on a weight basis), ten times the potency of affinity-purified CroFab antivenom on a weight basis, and seven times the potency of CroFab on a per molecule basis. The greater relative potency of the standalone FETUA protein combination is noteworthy, both for the small quantity of protein required to neutralize venom and because the FETUA combination lacks inhibitors of other types of venom toxins, which snake sera and antivenom contain. The potential for FETUA proteins to improve antivenom potency raises the question of their prospective polyvalency – their ability to inhibit MP activities of diverse species.

### *C. atrox* FETUA proteins inhibit the MP activities and lethality of evolutionarily distant pit viper venoms

The distinct, complementary, and combined contributions of the FETUA-2, FETUA-3, and FETUA- 5 proteins’ inhibition of *C. atrox* and *C. adamanteus* venom MPs, two well-diverged species, suggest that this set of proteins inhibits a spectrum of *Crotalus* MPs. Because snake venom MPs share structural features in their metalloproteinase domains (34, 48), and FETUA protein sequences are well-conserved among *Crotalus* species and other pit vipers (38) we reasoned that *C. atrox* FETUA proteins may be able to recognize and inhibit MPs of evolutionarily more distant species. To test this hypothesis, we surveyed the ability of *C. atrox* FETUA proteins to inhibit various MP activities of a selection of four pit viper (Crotalinae) venoms chosen on the basis of their increasing phylogenetic distance from *C. atrox* including: representatives from two other North American genera *Sistrurus tergeminus edwardsii* (divergence ∼ 8.5-12 mya) and *Agkistrodon contortrix* (divergence ∼13- 20 mya); and two Asian genera *Calloselasma rhodostoma* and *Deinagkistrodon acutus* (divergence ∼34 mya) (7, 45, 46, 49). We note that because *D. acutus* possesses orthologs of all five *Crotalus fetua* genes, and only these *fetua* genes, we infer that all five FETUA proteins in *Crotalus* were present in the most recent common ancestor of all pit vipers(38). Thus, the comparison of *C. atrox* FETUA-2-5 protein activities on these pit viper venoms addresses the conservation or divergence of FETUA-MP interactions across the timescale of the pit viper radiation.

Venom MP enzyme activities vary on different protein substrates. Thus, in order to assess the ability of both individual and combinations of FETUA proteins to inhibit a spectrum of MP activities from different pit viper venoms, we examined MP activity on a variety of biologically relevant protein substrates including collagen I, fibrinogen, and the basement membrane protein nidogen(50). For our initial test, we incubated increasing amounts of a mixture of all four evolutionarily derived *Crotalus* FETUA proteins (FETUA-2/3/4/5) with each whole venom and observed that this mixture of FETUAs was able to inhibit ≥95% of the type I collagenase activities of three divergent pit viper venoms (Fig. 4A; *A. contortrix* venom lacked significant collagenase activity and was omitted from this assay). Since titrations of the *C. atrox* FETUA proteins show that it does not require greater amounts of this mixture to strongly inhibit the evolutionarily more distant venoms, we infer that the strength of FETUA-MP protein interactions detected in this assay are similar across the venoms tested.

**Figure 4.**
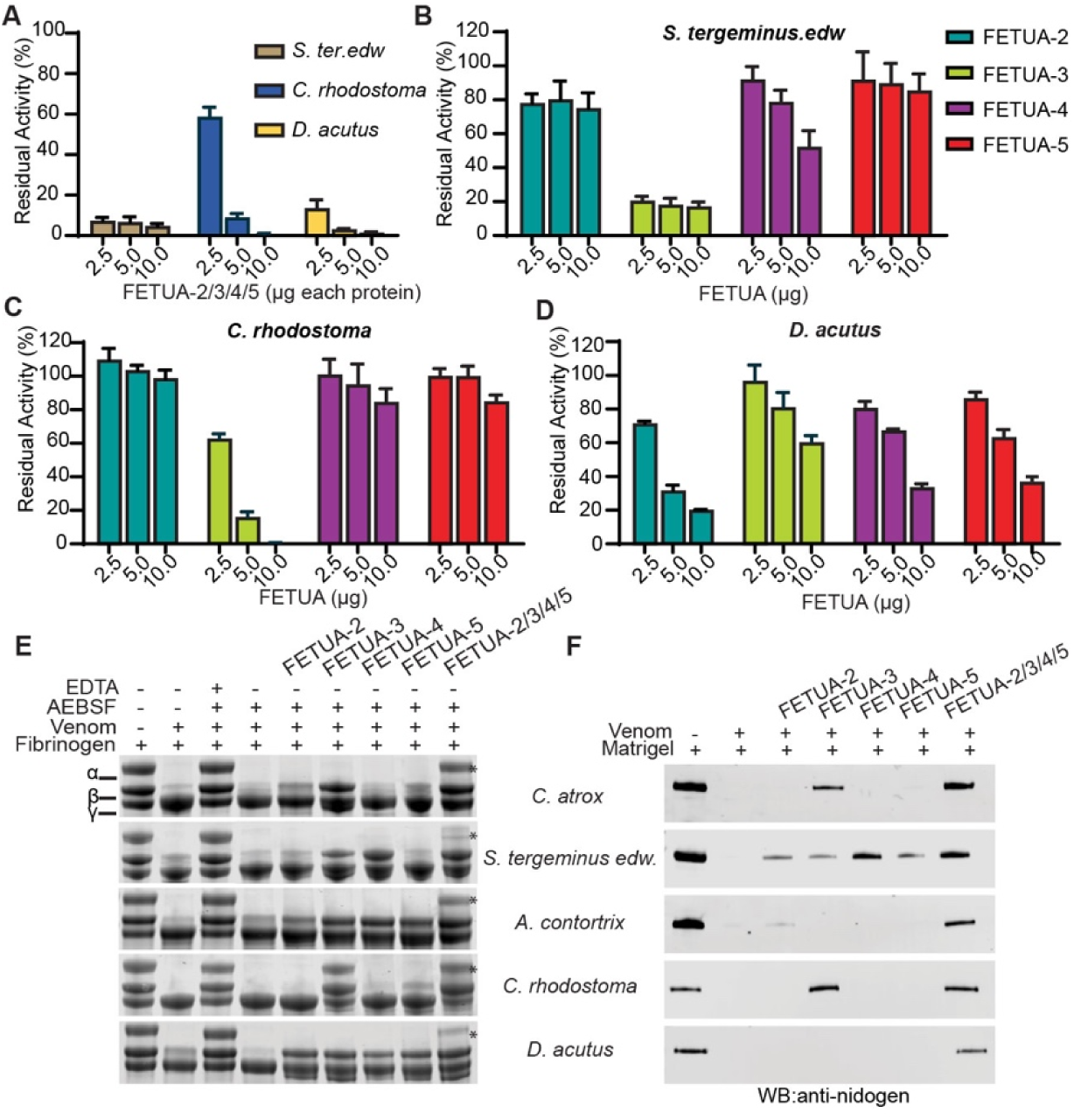
*C. atrox* FETUA proteins inhibit venom metalloproteinases in vitro from evolutionarily distant species. **A-D**) Inhibition of whole viperid venoms by individual and combinations of FETUA proteins. **A)** Inhibition of the whole venom collagenase activity of 10µg *S. tergeminus edwardsii*, 10µg *C. rhodostoma*, or 10µg *D. acutus* venoms by the combination of FETUA-2/3/4/5. The amount of each individual FETUA protein in the combination is shown on the X-axis and the fraction of residual MP activity is shown on the Y-axis. Note that all four venoms are strongly inhibited by combinations containing 5µg and 10µg of each FETUA protein. **B)** *S. tergeminus edwardsii* venom (10µg) activity is inhibited most strongly by FETUA-3 moderately by FETUA-2 and FETUA-4, and slightly by FETUA-5. **C)** *C. rhodostoma* venom (10µg) activity is strongly inhibited by FETUA-3, very weakly by FETUA-4 or FETUA-5, and not by FETUA-2. **D)** *D. acutus* venom (10µg) activity is inhibited most strongly by FETUA-2, and moderately by FETUA-3, FETUA-4, and FETUA-5. **E)** Inhibition of 1µg of whole venom MP fibrinogenase activities by individual (2 µg) and combinations of FETUA proteins (2 µg each). Each FETUA inhibits Bβ-fibrinogenase activity of multiple venoms, and combinations of FETUA proteins complement one another to inhibit A⍺-fibrinogenase activity in each venom (asterisks). MP fibrinogenase activity is revealed in lane 4 by the degradation of the ⍺- or β-chain of human fibrinogen by venom in the presence of 1mM AEBSF, a serine protease inhibitor, relative to the undigested control in lane 1. MP fibrinogenase activity is inhibited by the addition of 10mM EDTA. **F**) Inhibition of 1µg whole venom digestion of nidogen by individual (2µg) and combinations (2µg each) of FETUA proteins. Each venom digests nidogen as shown by the disappearance of the nidogen band in lane 2 relative to the undigested control in lane 1. The combination of FETUA-2/3/4/5 inhibits nidogen digestion by each venom, while individual FETUA proteins inhibit nidogen digestion in selected venoms.

To explore these interactions in greater detail, we tested the ability of individual *C. atrox* FETUA proteins to inhibit whole venom MP collagenase activity. The patterns observed revealed that all four FETUA proteins are able to inhibit MP activities in evolutionarily distant venoms, with each FETUA exhibiting some activity against *S. tergeminus* and *D. acutus* venoms (Fig. 4B and Fig. 4D), and FETUA-3 demonstrating very strong activity, and FETUA-4 and FETUA-5 slight activity, against *C. rhodostoma* venom in the collagenase assay (Fig. 4C).

Examination of FETUA inhibition of MP fibrinogenase activity also revealed broad cross-species FETUA-MP interactions. *C. atrox* and the four divergent species venoms cleave the A⍺- and Bβ- chains of fibrinogen in vitro. We show that these activities are due to MPs because they are inhibited by EDTA which blocks MP activity and not inhibited by the serine protease inhibitor 4-(2-aminoethyl)benzenesulfonyl fluoride (AEBSF) (Fig. 4E; compare lanes containing EDTA and AEBSF with those containing only AEBSF). Each individual FETUA protein inhibited Bβ-fibrinogenase activities in one or more venoms, with all four FETUAs exhibiting activity against *D. acutus* venom (Fig. 4E). Moreover, when all FETUAs were combined, A⍺-fibrinogenase activity was also inhibited to some degree in each of the four venoms (Fig. 4E, asterisks). This latter result suggests that different FETUAs inhibit different subsets of MPs in these venoms such that combining them inhibits a broader spectrum of MP activities.

We also found that the combination of all four FETUA proteins was able to inhibit the digestion of the basement membrane protein nidogen (Figure 4F) by *C. atrox* venom and all four divergent venoms. Several individual FETUA proteins also inhibited nidogen digestion by certain species venoms, including FETUA-3 which inhibited *C. atrox, S. tergeminus*, and *C. rhodostoma* venom, FETUA-2 which inhibited both *S. tergeminus* and *A. contortrix* venoms, and FETUA-4 and FETUA-5 which both inhibited *S. tergeminus* venom (Fig. 4F).

The in vitro assays demonstrate that individual *C. atrox* FETUA proteins recognize and inhibit MPs from evolutionarily distant pit vipers. With respect to their potential as components of antivenoms, a more demanding test is whether these FETUA proteins are able to neutralize the lethal action of diverse venoms in vivo. Therefore, we pre-incubated a combination of the FETUA-2/FETUA-3/FETUA-5 proteins with 2.5 x LD_50_ of the venoms of the two most evolutionarily divergent pit viper species, *C. rhodostoma* and *D. acutus*, and injected the mixtures intraperitoneally into mice. We observed that the *C. atrox* FETUA protein mixture fully protected mice from the lethal effects of *C. rhodostoma* venom (Figure 5A, P=.0027) and significantly protected mice from the lethal effects of *D. acutus* venom (Figure 5B, Table 4; P= 0.0004)(Table 5). The protection observed is particularly notable in light of the ∼34 million years of evolutionary divergence between these species and the *Crotalus* lineage and the different compositions of the venoms tested.

**Figure 5.**
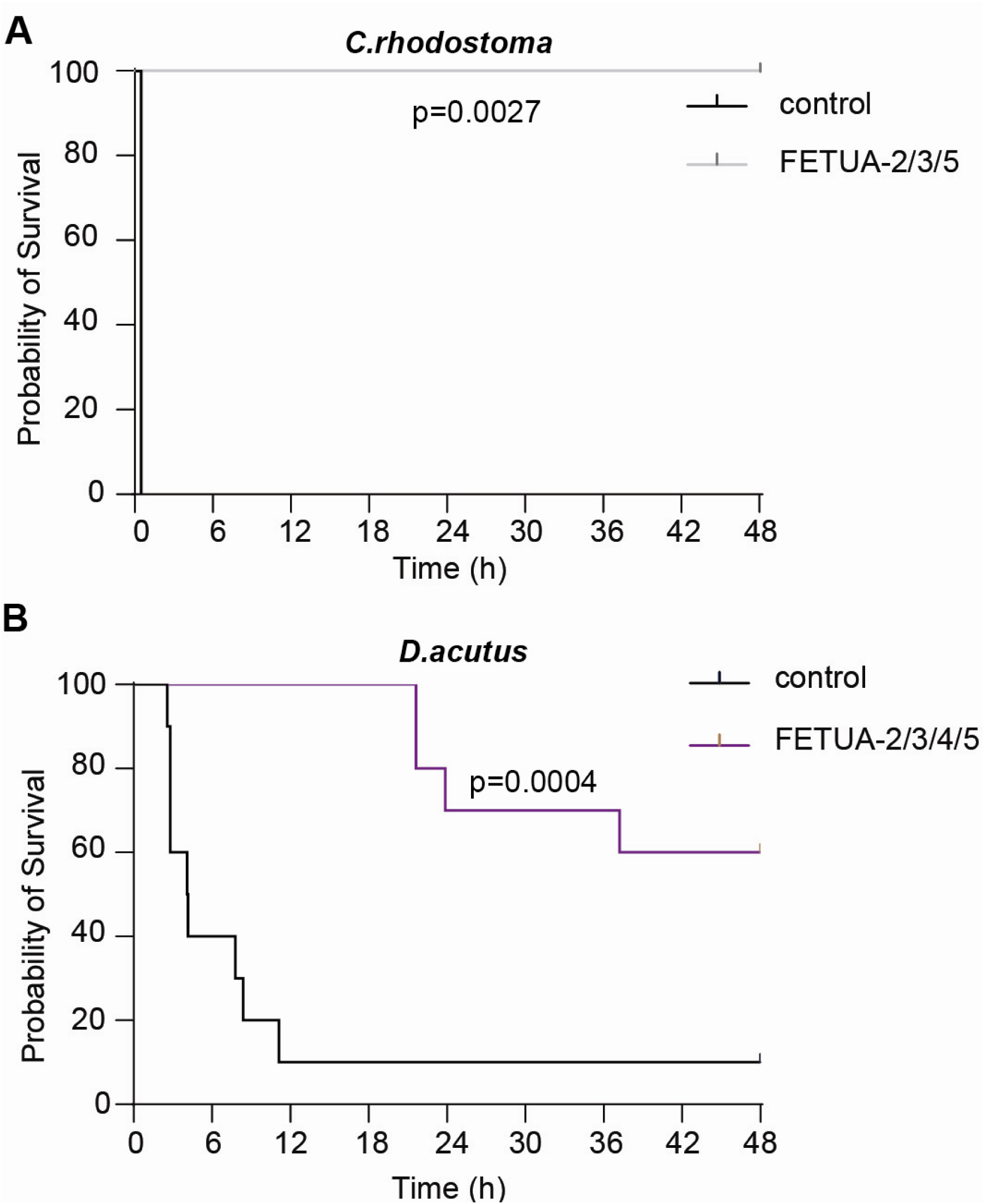
Combinations of *C. atrox* FETUA proteins inhibit the lethality of C. *rhodostoma* and *D. acutus* venoms. A. Kaplan-Meier survival plots of mice (n=5) that were injected i.p. with 2.5 x LD50 of *C. rhodostoma* venom that was preincubated with either saline (control) or the combination of 400 µg each of *C. atrox* FETUA-2, FETUA-3, and FETUA-5. and monitored for 48 hr. All mice survived with no apparent ill effects. B. Kaplan-Meier survival plots of mice (n=10) that were injected i.p. with 2.5 x LD50 of *D. acutus* venom that was preincubated with either saline (control) or the combination of 250 µg each of C. atrox FETUA-2, FETUA-3, FETUA-4, and FETUA-5. and monitored for 48 hr. The survival of one control mouse is interpreted to be due to mis-injection.

**Table 5.** Inhibition of Old World Pit Viper Venom Lethal Activities by Combinations of *C. atrox* FETUA Proteins.

|  | *C. rhodostoma* venom<sup>#</sup> | *D. acutus* venom* |
| --- | --- | --- |
| Treatment | Lethal Activity<br>(Live/Dead mice) | Lethal Activity<br>(Live/Dead mice) |
| Venom Control Only | 0/5 | 1/9 |
| FETUA-2, FETUA-3, FETUA-5 | 5/0 (P = 0.0027) | 6/4 (P=0.0004) |

#Mice were injected intraperitoneally with 2.5xLD50 of *C. rhodostoma* venom that had been pre-incubated with saline, or a mixture containing 400ug of each of the FETUA proteins. Animals were observed for a 48-hour observation period.
\* Mice were injected intraperitoneally with 2.5xLD50 of *D. acutus* venom that had been pre-incubated with buffer, or a mixture containing 250µg of each of the FETUA proteins. Animals were observed for a 48-hour observation period.

### FETUA- 2, FETUA-3, and FETUA-4/5 evolved deep within vipers

The abilities of the *C. atrox* FETUA proteins to inhibit the MP activities of evolutionarily diverse pit viper venoms in vitro and in vivo raises the question of whether these proteins might also act on MPs of more divergent venoms. Pit vipers (Crotalinae) are a subfamily of the family Viperidae which includes two other subfamilies, true vipers (Viperinae) among which are many medically important species found in Asia, Africa, and Europe, and Fea’s vipers (Azemiopinae), a small taxon that is a sister group to Crotalinae. We took two approaches to explore the potential ability of FETUA proteins to inhibit other viper venoms: a survey of *fetua* gene distribution and evolution, and an assessment of the ability of C*. atrox* FETUA proteins to inhibit MP activities in true viper (Viperinae) venoms.

The number and identity of *fetua* genes across the Viperidae family have not been examined previously. Fortunately, a significant number of viperid genomes representing key groups within the clade have recently become available. We annotated the *fetua* gene complex in these genomes (Fig. 6) and performed phylogenetic analyses of FETUA protein sequences to ascertain gene and protein identities (Fig. S3).

**Figure 6.**
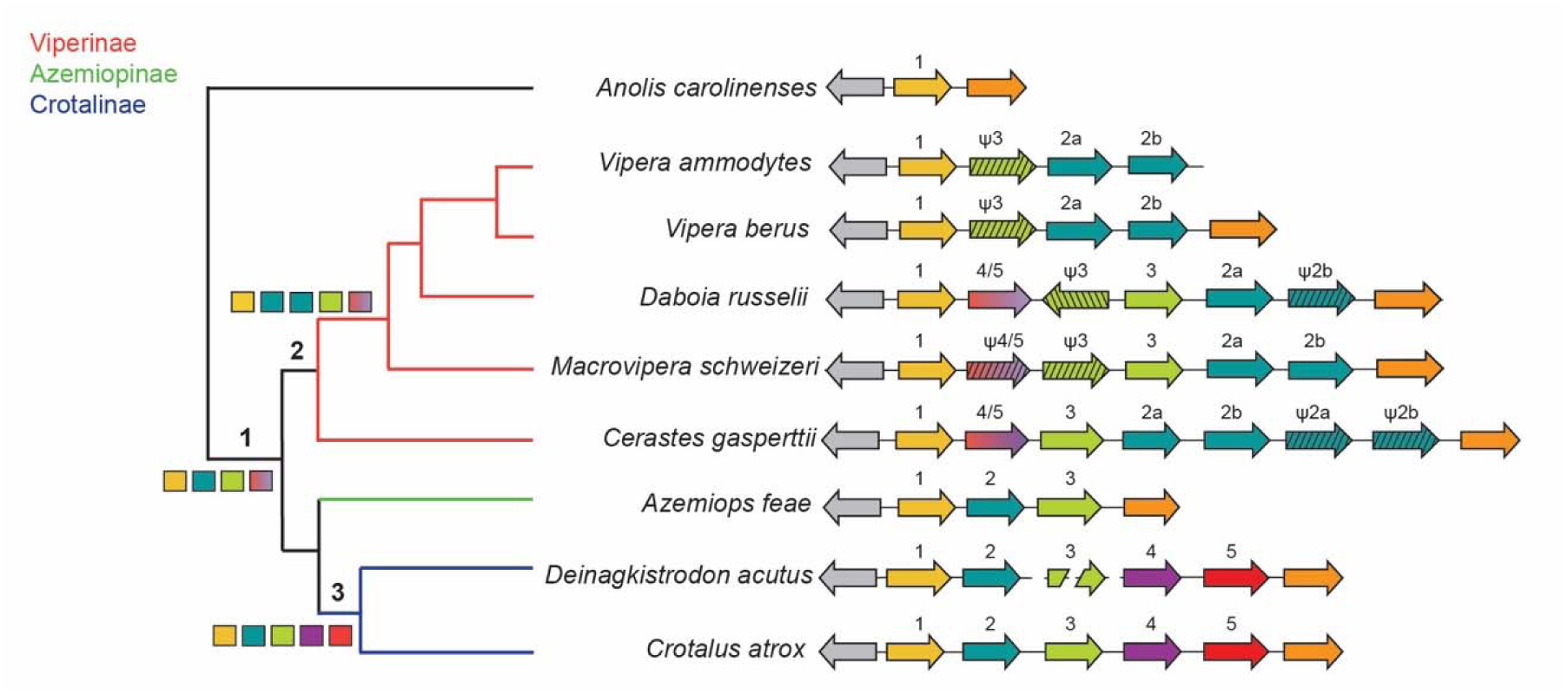
The FETUA-2, FETUA-3, and FETUA4/5 proteins evolved early within vipers. A schematic cladogram showing the number, identity, and arrangement of *fetua* genes in a sample of viper species belonging to the three subfamilies Crotalinae (blue), Azemiopinae (green), and Viperinae (red). Each fetua gene paralog is identified on the basis of gene phylogeny (fig. S3). The orientation of each locus is denoted by the direction of the arrow and pseudogenes are denoted by the Ψ symbol. The grey and orange arrows denote the position of the syntenic flanking genes *senp5* and *fetuin-B*, respectively. The presence of all five Crotalus *fetu*a orthologs in *D. acutus* supports the inference that all five genes were present in the most recent common ancestor of Crotalinae (node 3). The distribution of *fetua-1*, two *fetua-2* loci, *fetua-3*, and a homolog of *fetua-4/fetua-5* within Viperinae supports the inference that these genes were present in their most recent common ancestor (node 2). The loci shared between the most recent common ancestors of Viperinae and Crotalinae supports the inference that the most recent common ancestor of all vipers possesses *fetua-1, fetua-2, fetua*-3, and a homolog of *fetua-4/5* (node 1).

One key informative taxon is *Azimeops feae*, a member of the sister taxon to pit vipers that possesses just the three genes *fetua-1*, *fetua-2*, and *fetua-3* (Fig. 6; Fig. S4A, B). This observation indicates that the origins of *fetua-2* and *fetua-3* pre-date the radiation of the pit vipers. To trace the deeper evolutionary history of *fetua* genes, we examined the genomes of several true vipers (Viperinae), a clade that split from the pit vipers more than 40 million years ago (51). In several examined viper species’ genomes, we find only the *fetua-1*, *fetua-2,* and *fetua-3* genes, with the number and integrity of particular loci varying. Specifically, in *Macrovipera schweizeri* we find four intact *fetua* genes including one *fetua-1* locus, two *fetua-2* loci (we denote them *fetua-2a* and *fetua-2b*), and one *fetua-3* gene, while in *Vipera berus* and *V. ammodytes* we also identify one *fetua-1* locus, two *fetua-2* loci, and one *fetua-3* gene that has been pseudogenized by deletion mutations (Fig. 6). In *Daboia russelli* and *Cerastes gaspertii*, in addition to intact *fetua-*1, *fetua-2*, and *fetua-3* genes, we also find one homolog of the pit viper *fetua-4* and *fetua-5* genes (Fig. 6). The latter two loci in pit vipers are sister genes that encode proteins with similar sequences, we cannot assign a *fetua-4* or *fetua-5* identity to the viper paralog so we refer to this gene as *fetua-4/5* (Fig. 6; Fig. S4C).

Based upon the distribution of *fetua* genes and the phylogenetic relationships among these species, we infer that the FETUA-2 and FETUA-3 proteins and a homolog of the FETUA-4/FETUA-5 proteins existed in the last common ancestor of all viperids (Fig. 6, node 1). We also infer that specific genes were duplicated in certain lineages (*fetua-2* in true vipers and *fetua-4/5* in pit vipers) and that some *fetua* genes or proteins were lost in some lineages (FETUA-3 and FETUA-4/5 in *Vipera*; FETUA-4/5 in *Macrovipera* and *Azemiops*).

### *C. atrox* FETUA proteins inhibit the MPs and lethality of selected viper venoms

Since FETUA-2 and FETUA-3 have a long evolutionary history that pre-dates the radiation of the Viperidae and all viper FETUA proteins bear some relationship to and sequence homology with those found in pit vipers, we reasoned that *C. atrox* FETUA proteins might also inhibit true viper venom MPs. Therefore, we carried out similar in vitro tests as those above with two representative, medically significant true viper venoms *Echis carinatus sochureki* and *Bitis arietans*.

We found that a mixture of all four *C. atrox* FETUA proteins was able to inhibit ≥ 95% of *B. arietans* venom collagenase activity, and that all four individual FETUA proteins also exhibited some activity against *B. arietans* venom (Fig. 7A) (*E. carinatus* venom did not exhibit significant collagenase activity in this assay). We also observed inhibition of *B. arietans* venom Bβ-fibrinogenase activity by FETUA-3, FETUA-4, and FETUA-5, and inhibition of A⍺-fibrinogenase activity by the combination of all four FETUA proteins (Fig. 7B). *E. carinatus* venom exhibited somewhat weaker Bβ-fibrinogenase activity than *B. arietans*; this activity was inhibited by FETUA-2 and FETUA-3 and there was slight inhibition of venom A⍺-fibrinogenase activity by the combination of all four FETUA proteins (Fig. 7B). We also found that FETUA-3 strongly inhibited digestion of nidogen by *B. arietans* venom, and that FETUA-2 strongly inhibited digestion of nidogen by *E. carinatus* venom (Fig. 7C).

**Figure 7.**
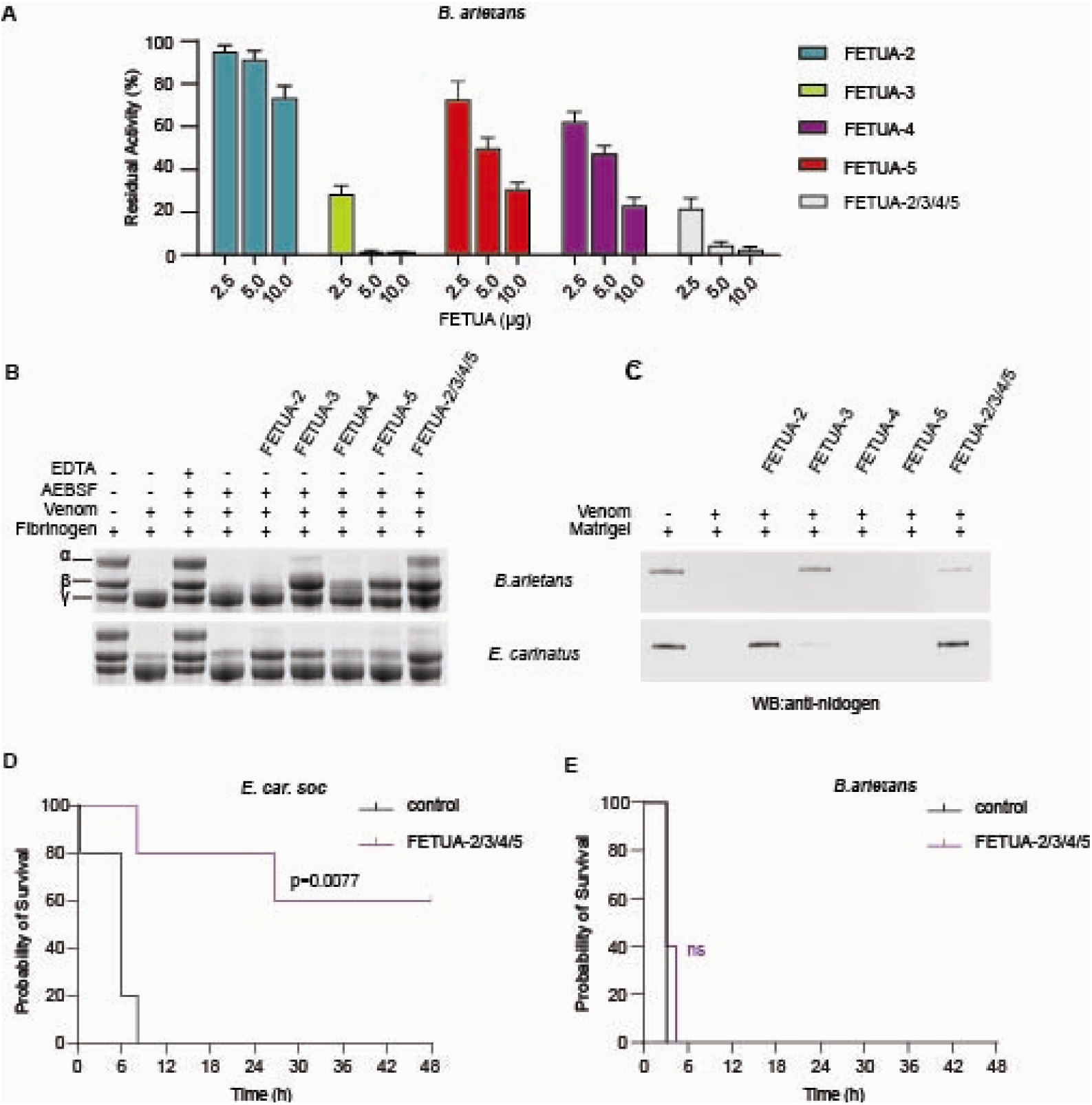
*C. atrox* FETUA proteins inhibit the metalloproteinases and lethality of selected viper venoms. **A**. Inhibition of *B. arietans* whole venom collagenase activity by individual and combinations of C. atrox FETUA proteins. Collagenase activity is strongly inhibited by FETUA-3 and FETUA-2/3/4/5,and moderately inhibited by FETUA-4, and FETUA-5 **B**. Selected *C. atrox* FETUAs inhibit viper whole venom fibrinogenase activity. Note that FETUA-3, FETUA-4, and FETUA-5 inhibit *B. arietans* B Bβ-fibrinogenase activity and that the combination of all four FETUA proteins inhibits A⍺-fibrinogenase activity. *C. atrox* FETUA 2- and FETUA-3 inhibit *E. carinatus* venom Bβ-fibrinogenase activity. **C**. *C. atrox* FETUA-3 and *C. atrox* FETUA-2 inhibit *B. arietans* and *E. carinatus venoms’ digestion of nidogen, respectively*. **D**. Kaplan-Meier survival plots of mice that were injected i.p. with 2.5 x LD_50_ of *E. carinatus sochureki* venom that was preincubated with either saline (control) or the combination of 60 µg each of *C. atrox* FETUA-2, FETUA-3, FETUA-4, and FETUA-5, and monitored for 48 hr. **E.** Kaplan-Meier survival plots of mice that were injected i.p. with 2.5 x LD_50_ of *Bitis arietans* venom that was preincubated with either saline (control) or the combination of 135 µg each of *C. atrox* FETUA-2, FETUA-3, FETUA-4, and FETUA-5, and monitored for 48 hr.

These in vitro assays reveal some activity of each *C. atrox* FETUA protein on one or both viper venoms’ MPs (i.e. polyvalency), despite ∼50 myr divergence between the lineages. We then tested the FETUA proteins’ ability to neutralize the lethal effects of each venom in vivo. We preincubated a combination of all four FETUA proteins with 2.5 x LD_50_ of each venom (in FETUA protein:venom protein weight ratios sufficient to inhibit in vitro activities) and injected the mixtures intraperitoneally into mice. We observed significant protection against the lethal effect of *E. carinatus sochureki* venom (Fig. 7D, P= 0.0077) but no protection against *B. arietans* venom (Table 6). The partial protection against *E. carinatus* venom suggests that the *C. atrox* FETUAs substantially inhibit important *E. carinatus* MP toxins. The lack of protection observed against *B. arietans*, despite the ability of the FETUAs to inhibit venom MP in vitro activities, may be due to insufficient inhibition of certain venom MPs by the heterologous *C. atrox* FETUAs and/or the action of non-MP venom toxins that are not impeded by the FETUA proteins.

**Table 6.** Inhibition of Viper Venom Lethal Activities by Combinations of *C. atrox* FETUA Proteins.

|  | *E. car. soch* venom<sup>#</sup> | *B. arietans* venom* |
| --- | --- | --- |
| Treatment | Lethal Activity<br>(Live/Dead mice) | Lethal Activity<br>(Live/Dead mice) |
| Venom Control Only | 0/5 | 0/5 |
| FETUA-2, FETUA-3, FETUA-4<br>FETUA-5 | 3/2 (P=0.0077 ) | 0/5 |

#Mice were injected intraperitoneally with 2.5xLD<sub>50</sub> of *E. carinatus sochureki* venom that had been pre-incubated with buffer, or a mixture containing 60µg of each of the FETUA proteins. Animals were observed for a 48-hour observation period.
\* Mice were injected intraperitoneally with 2.5xLD<sub>50</sub> of *B. arietans* venom that had been pre-incubated with saline, or a mixture containing 135µg of each of the FETUA proteins. Animals were observed for a 48-hour observation period.

## Discussion

This study was motivated by the long-established observations that many vipers are resistant to the lethal and other toxic effects of their own venoms, and that resistance can be transferred to experimental animals from snake sera. These phenomena prompt the hypotheses that these snakes might have “solved” the challenge of neutralizing many different toxins (or at least those most dangerous to themselves and other species) by co-evolving an array of specific toxin inhibitors, and that such inhibitors may be useful as antivenom components.

We have made three principal findings here that bear on these hypotheses. First, we discovered that specific combinations of *C. atrox* FETUA proteins complement one another’s MP-inhibiting activities and are sufficient to fully neutralize *C. atrox* venom’s lethal action in vivo with approximately ten times greater potency than affinity-purified commercial antivenom. Second, we show that these *C. atrox* FETUA proteins can inhibit the metalloproteinases and neutralize the lethality of other rattlesnakes and of evolutionarily distant pit viper and true viper venoms, demonstrating a high degree of polyvalency. And third, we found that true vipers possess similar sets of FETUA proteins as pit vipers, including both orthologs and closely-related paralogs of the FETUA proteins studied here. Our findings raise questions about how FETUAs inhibit such a broad array of MPs and prevent the lethality of hemorrhagic venoms, and they have implications for the potential use of recombinant MP inhibitors in formulating potent, polyvalent viper antivenoms.

### The distinct, complementary, and combined actions of FETUA proteins on venom MPs

Prior to this study, knowledge of FETUA protein functions was derived from three sets of observations regarding FETUA-2 and FETUA-3. First, it was well-established that the FETUA-2 proteins of the Asian pit vipers *P. flavoviridis* (HSF; (35) and *G. blomhoffi* (MSF)(36) and the South American pit viper *Bothrops jararaca* (BJ46a; (37) inhibit certain MPs and venom hemorrhagic activities. Second, it was reported that two FETUA-3 proteins from two Asian pit vipers P*. flavoviridis* and *G. blomhoffi* (named HLP and HLP-A, respectively) lack venom MP inhibitory activity (52). And third, we had found that, contrary to its reported lack of activity in Asian pit vipers, the *C. atrox* FETUA-3 protein is a strong inhibitor of MPs belonging to three different classes while *C. atrox* FETUA-2 had limited inhibitory activity to those MPs in vitro (38).

These observations led us to propose that FETUA-3 had acquired functions in the *Crotalus* lineage enabling it to become a major MP inhibitor. Therefore, we were initially surprised and puzzled by the lack of FETUA-3 activity in initial in vivo tests, which neither inhibited hemorrhage nor delayed lethality caused by *C. atrox* venom. This lack of activity prompted us to examine the ability of other individual FETUA proteins to inhibit hemorrhage and to discover that FETUA-2 and FETUA-5 inhibited hemorrhagic activity.

We infer from these results (and previous in vitro enzyme inhibition assays) that different individual FETUA proteins act on distinct subsets of MPs. The very strong anti-hemorrhagic activity of FETUA-2 and the partial activity of FETUA-5 indicate that these proteins act on the MPs primarily responsible for causing hemorrhage, and that FETUA- 3 (and FETUA-4) does not. A variety of individual *C. atrox* MPs have been isolated and found to have hemorrhagic activities spanning about three orders of magnitude, with MDC-6a (Atrolysin-a) the most potent hemorrhagic enzyme (MHD 0.04µg; (53) and MDC-4 (VAP-2) among the least hemorrhagic toxins (MHD 15µg;(54)). The low MHD (1.5µg) of whole *C. atrox* venom used in our studies leads us to infer that we are primarily observing the in vivo activity of the most potent hemorrhagic MP(s), of which FETUA-2 is a very strong inhibitor, FETUA-5 is a partial inhibitor, and FETUA-3 is a noninhibitor, whereas FETUA-3 is a strong inhibitor of MDC-4 and other select MPs collagenase activities in vitro(38).

Furthermore, several experiments here suggest that different FETUA proteins act in a complementary fashion. For example, we found that while FETUA-2 alone partially protected against the lethal activity of *C. atrox* venom, and FETUA-3 alone had no activity, the combination of the two proteins was fully sufficient to neutralize venom lethality. This suggests that FETUA-3 neutralizes important MP activities that FETUA-2 does not, and vice versa. Similarly, we found that while FETUA-2 and FETUA-5 each partially inhibit the hemorrhagic activity of *C. adamanteus* venom, the combination of the two proteins fully inhibited hemorrhage. One plausible interpretation of this result is that the two proteins act in a complementary fashion to inhibit different hemorrhage-causing *C. adamanteus* MPs, which can be tested by the isolation and identification of the respective MPs (the most potent of which appears to be distinct from *C. atrox* MDC-6a based on its reported molecular weight and glycosylation; (47).

In addition, the observation that the pairwise combinations of FETUA-2/FETUA-3 or FETUA-3/FETUA-5 do not neutralize *C. adamanteus* venom lethality, but that the combination of FETUA-2/FETUA-3/ FETUA-5 fully inhibits lethality indicates that the three FETUA proteins together inhibit a broader spectrum of *C. adamanteus* MPs than does each pair. The complementary activities of these FETUA proteins help to explain why a family of FETUA proteins has evolved and been maintained in Crotalus, i.e. for self-defense against a diverse array of venom MPs.

The ability of combinations of FETUA proteins to neutralize venom lethality and to prevent the appearance of envenomation symptoms in mice, in the absence of inhibitors of other types of venom toxins, is unexpected and unprecedented. Prior studies of FETUA-2 proteins from several species did not report the inhibition of venom lethality (35–37), or the lack thereof (we do not know whether this was because inhibition of venom lethality was not tested or not observed).

Our results raise the question of how blocking MP activities alone counteracts broader venom effects. The pathological effects of pit viper venom MPs are generally thought to arise from: i) their action on circulating proteins such as fibrinogen (as shown in Figure 4) which leads to coagulation defects(55); and ii) their breakdown of extracellular matrix (ECM) and basement membranes (BM), which compromises blood vessel walls and causes local and systemic hemorrhage as well as tissue and organ damage ((50). It has been postulated that MP action enables venom spreading and for toxins to gain access to the systemic circulation (via the lymph and blood)(56) such that they have been dubbed “gateway” toxins(57)Therefore, one possible explanation for how inhibition of MP activities prevents lethality in the mouse model here is that it limits the spread and access of other venom toxins to the general circulation. Further in vivo studies to track the spread of other toxins or to examine the ability of FETUAs to rescue animals after the administration of venom may shed additional light on how FETUAs prevent lethality.

### Evolutionary conservation and functional polyvalency of FETUA proteins

For FETUA proteins to be useful as antivenom components, they must be able to inhibit MP toxins from the array of species found in the target geographic area -- to exhibit polyvalency. We developed several lines of evidence that suggest that *C. atrox* FETUA proteins in particular and viperid FETUAs in general have polyvalent properties.

First, the ability of a combination of C. atrox FETUAs to neutralize *C. adamanteus* venom hemorrhagic and lethal activities directly demonstrates cross-species inhibition of MPs. Furthermore, since: i) *C. atrox* and *C. adamanteus* venoms contain representatives of most of the biochemical diversity of almost all known *Crotalus* MPs (15 of 16 paralog groups; (33, 44); ii) the time since their divergence from their most recent common ancestor (5.5-7 mya, (45, 46) is a substantial fraction of the overall divergence time of the *Crotalus* clade (7.5 -10 mya); and iii) the FETUA-2/3/4/5 proteins exhibit a high degree of sequence conservation across the clade(38), it appears reasonable to infer that the *C. atrox* FETUA proteins are polyvalent with respect to Crotalus species.

Second, we demonstrated the polyvalency of *C. atrox* FETUA proteins beyond Crotalus species by their individual and collective inhibition of MPs in vitro from a diverse array of viperid venoms including other North American pit vipers, Old World pit vipers (divergence ∼34 mya) and true vipers (divergence ∼50 mya) and by their neutralization of the lethal action of *C. rhodostoma*, *D. acutus,* and *E. carinatus* venoms in vivo. The ability of *C. atrox* FETUAs to inhibit MPs of distant species suggests that there is a conserved mechanistic basis for FETUA recognition and inhibition of MPs, which would bode well for the polyvalency of viperid FETUAs in general.

And third, our phylogenetic survey of the *fetua* gene family reveals that vipers possess a similar set of FETUA proteins as pit vipers, including orthologs of FETUA-2 and FETUA-3, and closely related paralogs of FETUA-2 and FETUA-4 and FETUA-5. Thus, the diversification of the MP toxin family throughout the evolution of viperids has unfolded in the continuing presence of a small set of structurally-related FETUA proteins.

The inability of the *C. atrox* FETUAs to neutralize *B. arietans* venom lethality in vivo, despite their inhibition of various venom MP activities in vitro, could be due incomplete inhibition of MPs in vivo and/or that blocking MP activity is insufficient to prevent lethality due to the action of other toxin types present in the venom. It will be valuable to test autologous (species-matched) viper FETUAs to assess their neutralizing activity.

### Nature’s antivenom: The prospect of potent, polyvalent antivenoms comprised of recombinant toxin inhibitors

Humanity has long relied on nature to provide medicines for our ailments. That dependency persists to the present with about half of all new pharmaceuticals developed over the past 50 years sourced from nature or derived from natural compounds including some of the most important first-in-class anti-infectives (ivermectin), anti-cancer agents (taxol), and cardiovascular drugs (lovastatin)(58). Many useful compounds are the products of chemical warfare among microbes or plants and their enemies (herbivores, pests). Driven by co-evolutionary processes, natural selection has tailored a pharmacopeia of molecules that frequently act as high-affinity, specific inhibitors of target enzymes, receptors, or ion channels.

Similarly, vipers have evolved specific toxin inhibitors as components of a self-defense system against their own venoms, and we have shown that the FETUA components of this system have co-existed and been co-evolving with MP toxins for approximately 50 million years. The practical question is whether these inhibitors possess characteristics that might improve upon the performance of current polyclonal antibody-based antivenoms derived from hyperimmune animals. We identify several properties of FETUA proteins that might endow them with clinically desirable properties of antivenom components. These include:

1. Specificity. The FETUA proteins bind to specific venom MPs with affinities in the nanomolar range (59); Dowell et al., unpublished), which are comparable to the affinities of polyclonal antibodies from hyperimmunized animals.
2. Mode of action. The FETUA proteins directly inhibit MP enzyme activity, and we have recently shown that FETUA-3 binds within the active site of multiple target enzymes (Dowell et al.,unpublished]. Antibody binding does not necessarily inhibit MP activity or target the enzymes’ active site.
3. Multi-target Inhibitors. FETUA-2 and FETUA-3 have each been demonstrated to inhibit multiple MPs from different paralog groups and classes(38, 60, 61). Thus, a small combination of FETUAs achieve a broad-spectrum of MP inhibition that polyclonal antivenoms may require many different antibody molecules to effect.
4. Potency. We have shown that it requires about one-tenth as much FETUA protein as affinity-purified CroFab antivenom protein on a weight basis to neutralize the lethality of *C. atrox* venom. The greater potency of the FETUA protein combination probably reflects a greater efficiency of the FETUA mixture in inhibiting MP activity due to its mode of action and inhibition of multiple targets. The affinity-purified CroFab antivenom contains antibodies to many other proteins that may not contribute to *C. atrox* venom neutralization. Since all other commercial antivenoms are not affinity-purified and consist mostly of proteins that are not venom-reactive, the potential relative potency of FETUA mixtures compared to other viper antivenoms may be much greater.
5. Species Polyvalency. The cross-species inhibition of MPs by FETUA proteins indicates that there is a significant degree of conservation of FETUA-MP protein-interaction interfaces over large evolutionary distances. This is likely to reflect some constraint on these interaction sites imposed by the need for individual FETUA proteins to inhibit multiple MP paralogs within individual species.

The polyvalency of antivenom antibodies depends on the degree of conservation of epitopes on proteins from species used to immunize animals with those of non-immunizing target species, which decays over time and with species divergence (62). By acting on more constrained sites on multiple targets, single FETUA proteins or a small combination of FETUAs appear to attain a degree of a polyvalency (e.g. pan-pit vipers or pan-vipers) that appears difficult for polyclonal antivenoms to attain.

There are several major unknowns that need to be addressed in considering FETUAs as prospective antivenom components. The first is which FETUAs to use for a given set of target species. We explored the use of only *C. atrox* FETUAs here and pursued experiments on increasingly distant viper venoms because we observed extensive cross-inhibition in vitro and in vivo. Optimal FETUA mixtures for a given set of target species may involve the use of proteins from those or closely related species. The second unknown is which other components are necessary or desirable to use in conjunction with FETUAs to control other viper toxins. We do not expect inhibition of MPs to be sufficient to prevent the lethality or control the morbidity of all viperid venoms due to the widely varying content, amounts, and potency of other toxins present. For example, there is abundant evidence that PLA_2_-derived toxins contribute greatly to viperid venom toxicity (12, 14, 16, 17), and therefore may need to be neutralized in conjunction with MPs, and perhaps other toxins as well. Third, the studies here all involved pre-incubation of the FETUAs with venoms. It will be important to test the FETUA mixtures in “rescue’ experiments by administering them separately after venom to determine the degree to which they can prevent mortality and morbidities once MP action has begun. A fourth unknown concerns FETUA proteins’ safety and pharmacokinetic behavior. FETUAs are naturally-occurring glycoprotein homologs of fetuin-A, the most abundant protein in fetal blood and a major serum component in all amniotes with a serum half-life on the order of days (63), and they are similar in size to antibody Fab fragments (∼50kDa), so they may possess some favorable pharmacological characteristics.

It is also not known how FETUA-containing antivenoms might compare with the new antibody and small molecule strategies that are currently being explored. Regardless of composition, antivenoms must neutralize the spectrum of major toxins in a set of target species. Several broadly cross-reactive monoclonal antibodies or nanobodies against elapid toxins have recently been described, combinations of which neutralize the venoms of a wide variety of elapid species (18–21) Whether the strategies used for the elapid toxins would also succeed for such diverse viper toxin families as the metalloproteinases, and yield antibodies with properties comparable to FETUAs is yet to be determined.

Small molecules such as the active site inhibitor marimastat have been shown to inhibit a broad spectrum of MP activities in a wide array of species (64) and to neutralize a variety of viper venoms when administered in conjunction with the PLA_2_ inhibitor varespladib in experimental animals (12, 14–17). This breadth of action and simple composition are very promising, but it is also to be expected that small molecules will have significantly different pharmacokinetic behaviors and off-target effects than antibodies or FETUAs (e.g. marimastat and varespladib also inhibit host enzymes). There is no clinical knowledge yet of these agents’ relative advantages or disadvantages in the management of snakebite.

Ever since the discoveries of venom resistance in snakes and of serum-borne toxin inhibitors, there has been conjecture about their prospects as antivenoms or antivenom components, but this possibility has not been realized. One key factor restraining that potential has been the incomplete knowledge of the viper self-defense system. We suggest that this new understanding of the efficacy and polyvalency of FETUA protein mixtures warrants further investigation of their prospective clinical use.

## Materials and Methods

### Recombinant protein production

All constructs and recombinant proteins were prepared commercially (GenScript, Piscataway, NJ, USA). The sequences encoding *C. atrox* FETUA proteins were based on sequences obtained from the genome of a *C. atrox* specimen (s238). To facilitate their expression as recombinant proteins, the endogenous signal peptide sequence (1-19 amino acids) was replaced with a sequence optimized for protein expression (MGWSCIILFLVATATGVHS) in mammalian cell lines. The synthetic genes were cloned into the mammalian expression vector pcDNA3.4 (Invitrogen, Carlsbad, CA, USA).

The recombinant FETUA protein expression plasmids were transiently transfected into CHO-S cells and grown in serum-free media to facilitate the isolation of the tag-free proteins. After five days, the 40-100 ml culture supernatants were harvested and the proteins (isoelectric points approximately 5.6-5.8) were isolated by applying the supernatants to a HiTrap Q Fast Flow anion-exchange column (Cytiva) that was equilibrated with a low salt loading buffer (20mM Tris pH 8.0, 0% NaCl (w/v)) and eluting with a linear salt gradient (20mM Tris pH 8.0, 0-30% NaCl (w/v). The purity of the recombinant proteins was analyzed by size-exclusion chromatography on a TSKgelG3000SWXL HPLC column developed with 0.1M Na_2_SO_4_ in 0.118M phosphate buffer pH 7.0 and estimated to be 95-99%. The samples were dialyzed against either Tris-buffered saline (TBS) or phosphate-buffered saline (PBS) and stored at -80°C.

Animals and biological materials.

All animals used in this work were housed at and in the care of the Viper Resource Center of the National Natural Toxins Research Center (NNTRC, Texas A&M University-Kingsville. Venom was obtained from *C. atrox*, *C. adamanteus*, *Sistrurus tergeminus edwardsii* (Texas; NNTRC specimen SCE-P, lot# 02112019)), and *Calloselasma rhodostoma* (Thailand; NNTRC specimen CR777, lot #02112019) specimens and blood *from C. atrox* individuals according to study protocols reviewed and approved by the Texas A&M Kingsville Institutional Animal Care and Use Committee in compliance with all applicable federal regulations governing the protection of research animals (Viper Resource Center at Texas A&M University-Kingsville, IACUC #:2024-12-2020).

Pools from 100 individuals of *C. atrox* (animals from Texas, New Mexico, and Arizona) and 100 individuals of *C. adamanteus* (all animals from Florida) were used in all assays, and the same venom pools were used for all in vivo experiments. *D. acutus* (pool from two individuals, lot #0325), *E. carinatus sochureki* (Pakistan origin, captive bred, pool from 12 individuals, lot # 0124) and *B. arietans* (Togo and Benin origin, pool from 12 individuals, lot #0125) venoms were purchased from the Kentucky Reptile Zoo. Venom stock solutions were made by dissolving venoms at 10 mg/mL in 20 mM Tris pH 8.0, 1 mM CaCl_2._ The solutions were filtered through a 0.2 µm Durapore syringe filter and stored at -80°C until use.

### Animal studies

Various standard assays for venom or antivenom activity necessitate the use of laboratory mice (40). The authors and the NNTRC have conducted this research in accordance with the 3R principles – Replacement, Reduction, and Refinement, by using in vitro enzyme assays where applicable, designing the scope and scale of animal experiments so as to minimize the number of animals required to attain meaningful and reliable results.

All animal procedures were conducted at the Viper Resource Center of the National Natural Toxins Research Center (NNTRC), Texas A&M University–Kingsville. All protocols were reviewed and approved by the Texas A&M University-Kingsville Institutional Animal Care and Use Committee (IACUC #: 2024-12-2020) and complied with all applicable federal regulations governing the care and use of laboratory animals.

BALB/c mice (male and female, 18–20 g) were used for all in vivo experiments. Animals were housed in Alternative Design Unicage systems (Model RC71U; 9 × 18 × 8 in.) with woodchip bedding at a controlled ambient temperature of 68°F under a standard 12 h light/dark cycle, with ad libitum access to food and water. Mice were housed in groups of 4–5 per cage and acclimatized 5 days prior to experimentation.

Animals were randomly assigned to experimental groups, with equal numbers of males and females. Humane endpoints included severe lethargy, immobility, or signs of systemic distress; however, in lethality studies, animals were monitored until experimental endpoints (up to 48 h) to allow accurate determination of survival curves, as approved by the institutional ethics committee. At the endpoints, the health status of each surviving mouse was monitored using the mouse grimace scale (65). Analgesics were not administered.

### Hemorrhage activity assay

A hemorrhage assay was used to determine the minimal hemorrhagic dose (MHD) of crude *C. atrox* and *C. adamanteus* venoms (40). A starting concentration of 0.2 mg/mL of each venom was prepared in 0.85% NaCl and a series of two-fold dilutions was made to obtain an additional four concentrations, of which 0.1 mL of each dilution was injected subcutaneously into the ventral side of BALB/c mice (male or female, 18-20g) to test for activity. After 24 hr, the mice were cervically sacrificed and the skin removed. The hemorrhagic spots were measured with a caliper, and the MHD was defined as the amount of venom protein required to produce a 10 mm^2^ hemorrhagic spot. This was determined to be 1.5µg for both species’ venoms.

### Antihemorrhagic assay

To test for the anti-hemorrhagic activity of individual FETUA proteins, 1 MHD of either *C. atrox* or *C. adamanteus* venom was pre-incubated with 10µg of a single FETUA protein or 10µg each of two FETUA proteins, in 0.1 ml of saline solution for 30 minutes at 37 °C. 0.1 mL of each venom/FETUA mixture (containing one MHD of venom) was injected subcutaneously into the ventral side of BALB/c mice (male or female, 18-20g) to test for activity. After 24 hr, the mice were cervically sacrificed, and the skin was removed. The animals were photographed and the area and intensity of the hemorrhagic spots were quantified using an AI hemorrhagic tool (66).

### LD_50_ determination

Standard antivenom studies test the ability of candidate antivenoms to neutralize an amount of venom that is a multiple of the venom LD_50_, in which the venom or antivenom/venom mixture is delivered by the intraperitoneal (i.p.) or the intravenous (i.v.) route. We selected the intraperitoneal route for our studies because prior experience showed that death occurred more slowly when venom was delivered by the i.p. than by the i.v. route (typically about two hours versus 5-10 minutes, respectively). We infer that this longer time to death reflects the time required for the venom to spread and cause systemic effects, thereby simulating natural envenomation to some degree.

To determine the LD_50_ for each venom, five groups of five mice (male and female BALB/c, 18-20 g) for each venom were housed in cages and observed throughout the quarantine period and experiments. Venoms were dissolved in physiological saline at the highest concentration of venoms that were used for injection. All solutions during the experiment were stored at 4 °C and warmed to 37 °C just before injection into mice. The lethal toxicity was determined by injecting 0.2 ml of venom (at various concentrations) into the peritoneum. The injections were administered using a 1 ml syringe fitted with a 30-gauge, 0.5-in. needle. Saline controls were used. The endpoint of lethality of the mice was determined after 48 h. The LD_50_ calculations were based on the Spearman and Karber method (Spearman C, 1978). The LD_50_ of the respective venoms were: *C. atrox*, 4.1mg/kg; *C. adamanteus*, 5.0 mg/kg; *C. rhodostoma*, 13.75 mg/kg; *D. acutus* 4.8mg/kg; *E. carinatus sochureki*, 1.2mg/kg; *B. arietans*, 2.7 mg/kg.

### ED_50_ determination

To determine the median effective doses (ED_50_) of *C. atrox* serum, CroFab antivenom, and *C. atrox* FETUA protein mixtures against *C. atrox* venom, five groups of five mice (male and female BALB/c, 18-20 g) were challenged with a mixture of three LD_50_ of C. atrox venom and varying amounts of serum, antivenom, or recombinant FETUA proteins in a standard antivenom assay (40). CroFab was reconstituted according to the manufacturer’s instructions. *C. atrox* serum and FETUA-protein mixtures were diluted in sterile saline, and two-fold serial dilutions were made with 0.85% sodium chloride to obtain four additional concentrations. A stock *C. atrox* venom solution containing 30 LD_50_s was freshly prepared at 0 °C on the day of the experiment. Venom and sera/antivenom/FETUA proteins were mixed together at a 1:2 volume ratio and incubated at 37 °C for 30 min before injections. Each mouse was injected with 0.2 mL of venom/antivenom mixture into the peritoneum. The mice were observed and the number of mice alive or dead was recorded at the 48 hr endpoint. ED_50_ was calculated using a Probit model (68).

### Lethality protection tests

To determine whether individual or combinations of *C. atrox* FETUA proteins were able to neutralize the lethal action of venoms, five mice (male and female BALB/c, 18-20 g) were challenged with a mixture of 3 x LD_50_ of *C. atrox* venom, 3 x LD_50_ of *C. adamanteus* venom, or 2.5 x LD_50_ of *C. rhodostoma*, *D. acutus*, *E. carinatus sochureki*, and *B. arietans* venoms and varying amounts of recombinant FETUA proteins in a standard antivenom assay (40). FETUA-protein mixtures were diluted in 1x PBS, stock venom solutions containing 17.5-21 LD_50_s were freshly prepared at 0 °C on the day of the experiment. Venom and FETUA proteins were mixed together at a ratio of 1:1.4 volume ratio and incubated at 37 °C for 30 min before injections. Each mouse was injected with 0.2 mL of venom/FETUA mixture into the peritoneum. The mice were observed for 48 h and their condition, time of death, and survival were recorded. Kaplan-Meier survival plots were drawn in GraphPad Prism version 10.6.1 and the significance of survival differences between groups was assessed using the log-rank (Mantel-Cox) test.

### Antivenom

CroFab (lot BN201894; expiration date May 2023), was obtained from ASD Healthcare and kept refrigerated as a lyophilized powder (note that antivenoms can retain their neutralization efficacy long beyond their expiration date; (69).

### Protein assay

The protein concentration of sera was determined by Bradford assay (70). FETUA and CroFab FAb protein concentrations were determined by measuring A_280_ in a Nanodrop microspectrophotometer using extinction coefficients calculated from the primary amino acid sequence of the mature FETUA proteins and IgG.

### SDS-PAGE

Protein samples were resolved by SDS-PAGE using 4-20% Criterion TGX™ Precast Midi Protein Gel for 40 minutes at 200 V (Criterion Cell; Bio-Rad). Samples were prepared in Laemmli buffer with or without 2-mercaptoethanol to achieve reducing or non-reducing conditions, respectively, and heated for 10 min at 95°C.

### Collagenase activity

Azocoll substrate (Sigma-Aldrich) was suspended at 5 mg/ml in 20 mM Tris-HCl pH 8.0, 2mM CaCl_2_. Working dilutions of venom were prepared from the venom stock solution in 20 mM Tris-HCl pH 8.0, 2mM CaCl_2_ at 1 mg/ml. Ten micrograms of venom was incubated with increasing amounts of inhibitor candidates in 50 μl microliters and incubated at 37 °C for one hour. *D. acutus* venom was preincubated with 2.5mM AEBSF (final) to inhibit serine protease activity before incubation with inhibitor candidates. The substrate (950 μl) was added to the enzyme-inhibitor solution and incubated at 37 °C for 60 minutes with agitation. Following incubation, samples were centrifuged at 15,000 x g (or RPM) for 2 min, the supernatant was collected, and absorbance measured at 520 nm. All assays were performed in triplicate.

### Fibrinogenolytic assay

One microgram of venom was preincubated with protease inhibitors (1 mM AEBSF and/or 10 mM EDTA, final concentrations) for 15 min at room temperature in PBS. Subsequently, the venom was incubated with FETUA proteins (2 µg of each protein) at 37 °C for 30 min, the 50 µg of human fibrinogen (50 µg; Enzyme research laboratories) was added in a final reaction volume of 20 µl, and the enzyme–inhibitor mixture incubated with shaking at 37 °C for 1 h. Samples were then loaded onto 8-16% gradient SDS–PAGE gels and analyzed by Coomassie Blue staining.

### Nidogen digestion assay

One microgram of venom was preincubated with protease inhibitors (1 mM AEBSF, final concentration) for 15 min at room temperature. Subsequently, the venom was incubated with FETUA proteins (2 µg of each protein) at 37 °C for 30 min, then 50 µg of Matrigel basement membrane extract (Corning) was added in a final reaction volume of 50 µl Tris-buffered saline, 2mM CaCl_2_ , and the enzyme–inhibitor mixture was incubated with shaking at 37 °C for 1 h. Samples were then loaded onto 4-20% gradient SDS–PAGE gels, transferred onto PVDF membrane, and analyzed by Western blotting using an affinity purified rabbit anti-human nidogen 1 antibody diluted 1:6000 (proteintech, 13766-1-AP) and an Alexa Fluor 680 tagged goat anti-rabbit IgG.

### Serine protease activity Assay

*C. atrox* venom serine protease activity was determined using N_α_-Benzoyl-L-arginine 4-nitroanilide hydrochloride (BapNA) chromogenic substrate. To measure the effect of the FETA proteins on venom serine proteases activity, 50 μg of venom was incubated with either 0 μg or 50 μg of individual FETUA proteins in 50mM Tris-HCl, 150mM NaCl, 10 mM EDTA, pH 7.8, in a total volume of 180 μl at 37°C for 30mins, in the wells of a 96- well microtiter plate. The reaction was initiated by the addition of BapNA to a final concentration of 0. 5mM. Hydrolysis of BapNA was monitored by measuring absorbance at 405 nm every 15 secs for 4 mins at 37°C. The rates of the catalyzed reactions were determined from the slopes of the linear plots of absorbance versus time, and serine protease activity was expressed as change in absorbance at 405 nm per minute.

### Phospholipase A_2_ activity assay

*C. atrox* venom PLA_2_ activity assay was performed using a phospholipase A2 activity kit (Abcam). To evaluate the effect of the FETUA proteins on venom PLA_2_ activity, 80 ng of venom was incubated with either 0 ng or 80 ng of individual FETUA proteins in PBS in a total volume of 40 μl at 37°C for 30mins. Following incubation,10 μl aliquots of each incubation mixture were added to corresponding wells of a second 96-well microtiter plate containing 5 μl assay buffer (25 mM Tris-HCl, 10 mM CaCl_2_,100 mM KCl, 0.3 mM Triton X-100, pH 7.5) and 10 μl of 10 mM 5,5-dithio-bis-(-2-nitobenzoic acid) (DNTB). The reaction was initiated by adding 200 μl of substrate solution (1.66 mM diheptanoyl thio-PC dissolved in assay buffer). Activity of PLA_2_ was monitored by measuring absorbance at 414 nm every 15 secs for 4 mins at 25°C. The rates of the catalyzed reactions were calculated from the slopes of the linear plots of absorbance versus time, and PLA_2_ activity was expressed as change in absorbance at 414 nm per minute.

### Phylogenetic analysis

Sequences used in our protein phylogenies were deduced from the hypothetical translation of assembled genomic regions from *D. russelli* or genomic regions from NCBI databases (see the full list of sequences Data S1). For phylogenetic analyses, we employed sequences from the first six exons because of the variable presence/absence of a deletion in the seventh exon of FETUA-2 and FETUA-3. To construct phylogenies, we aligned coding sequences using MAFFT v7.505 (71) with the automatic algorithm selection option (--auto), and their translated protein sequences were aligned using the global pairwise alignment strategy with iterative refinement (--globalpair --maxiterate 1000). IQ-TREE2 (72) was used to generate maximum likelihood phylogenetic trees from coding sequences and protein alignments with automatic model selection. Branch support was evaluated using 1000 ultrafast bootstrap replicates and 1000 SH-aLRT replicates.

## Acknowledgments

We thank Noah Dowell and Jory van Thiel for comments on the manuscript and Juan Salinas, Emelyn Salazar, and Andrea Esqueda of the National Natural Toxins Research Center for their venom extractions, assistance, and animal husbandry. This work was funded by the Howard Hughes Medical Institute (S.B.C.), the Andrew and Mary Balo and Nicholas and Susan Simon Endowed Chair at the University of Maryland (S.B.C), and the Viper Resource Center grant #P40OD01960-22 (NNTRC, Texas A&M University-Kingsville, E.E.S.)

