## Supplemental Information for "Nature’s Antivenom: Combinations of conserved rattlesnake serum metalloproteinase inhibitors block the lethal action of viper venoms"

Corresponding author Sean B Carroll

**This PDF file includes:**

Figures S1 to S4

**Other supporting materials for this manuscript include the following:**

Datasets S1

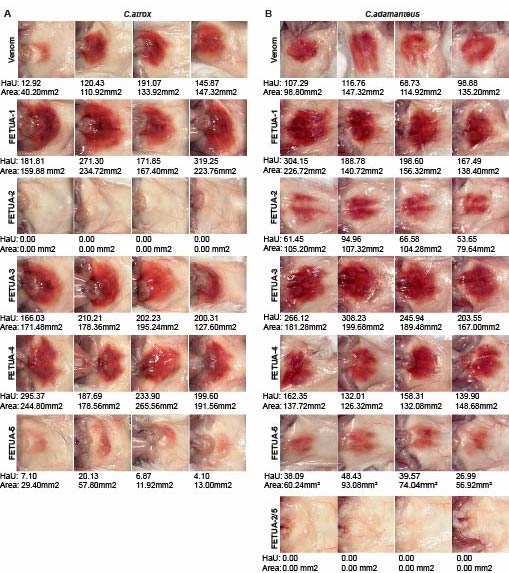

Fig. S1 Inhibition of rattlesnake venom hemorrhagic activity by selected FETUA proteins.

Groups of mice (n=4 per group) were injected subcutaneously with 1.5 µg of either (A) *C. atrox* venom or (B) *adamanteus* venom that was pre-incubated with either saline (venom control) or 10µg of individual or combinations of FETUA proteins indicated. The animals were sacrificed after 24 hr, the hemorrhage was photographed and hemorrhagic units (HaU) quantified by their area and intensity.

A**.** *C. atrox* venom hemorrhagic activity is completely inhibited by FETUA-2 and strongly inhibited by FETUA-5. FETUA-1, FETUA-3, and FETUA-4 enhance hemorrhage at the high protein concentration tested. The inhibition/enhancement calculations are tabulated in Table 1.

B. *C. adamanteus* venom hemorrhagic activity is partially inhibited by FETUA-2 and FETUA-5, and completely inhibited by the combination of FETUA-2 and FETUA-5.

FETUA-1, FETUA-3, and FETUA-4 enhance hemorrhage at the high protein concentration tested. The inhibition/enhancement calculations are tabulated in Table 3.

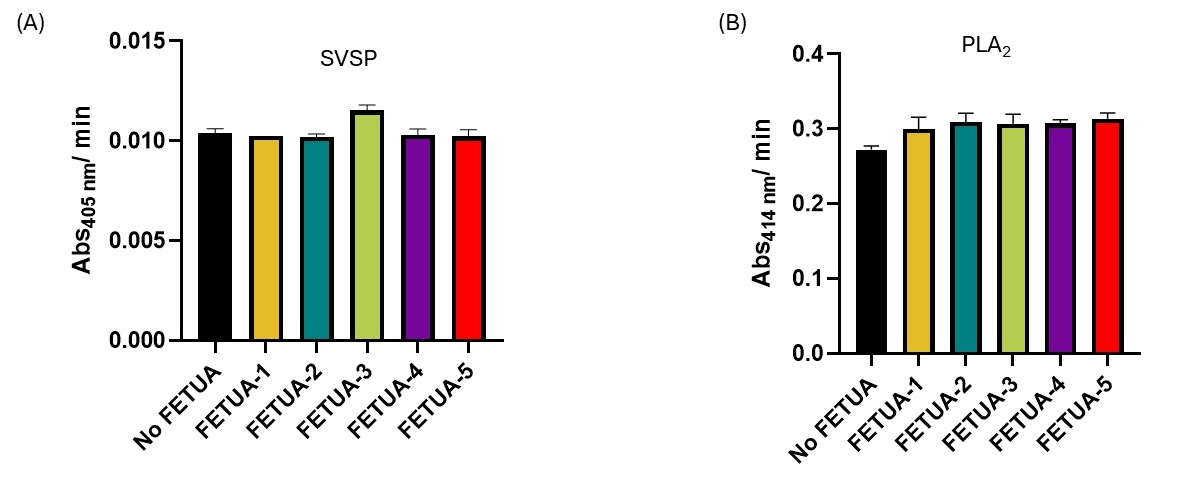

Fig. S2 FETUA proteins do not inhibit *C. atrox* venom serine protease (SVSP) and Phospholipase A2 activities.

(A) Bar plot showing the rate of hydrolysis of BapNA by *C. atrox* venom serine proteases in the presence of the indicated FETUA proteins.

(B) Bar plot showing the rate of hydrolysis of diheptanoyl thio-PC by of *C. atrox* venom PLA2 in the presence of the indicated FETUA proteins. Errors bars represent standard deviation of duplicate measurements from two separate assays.

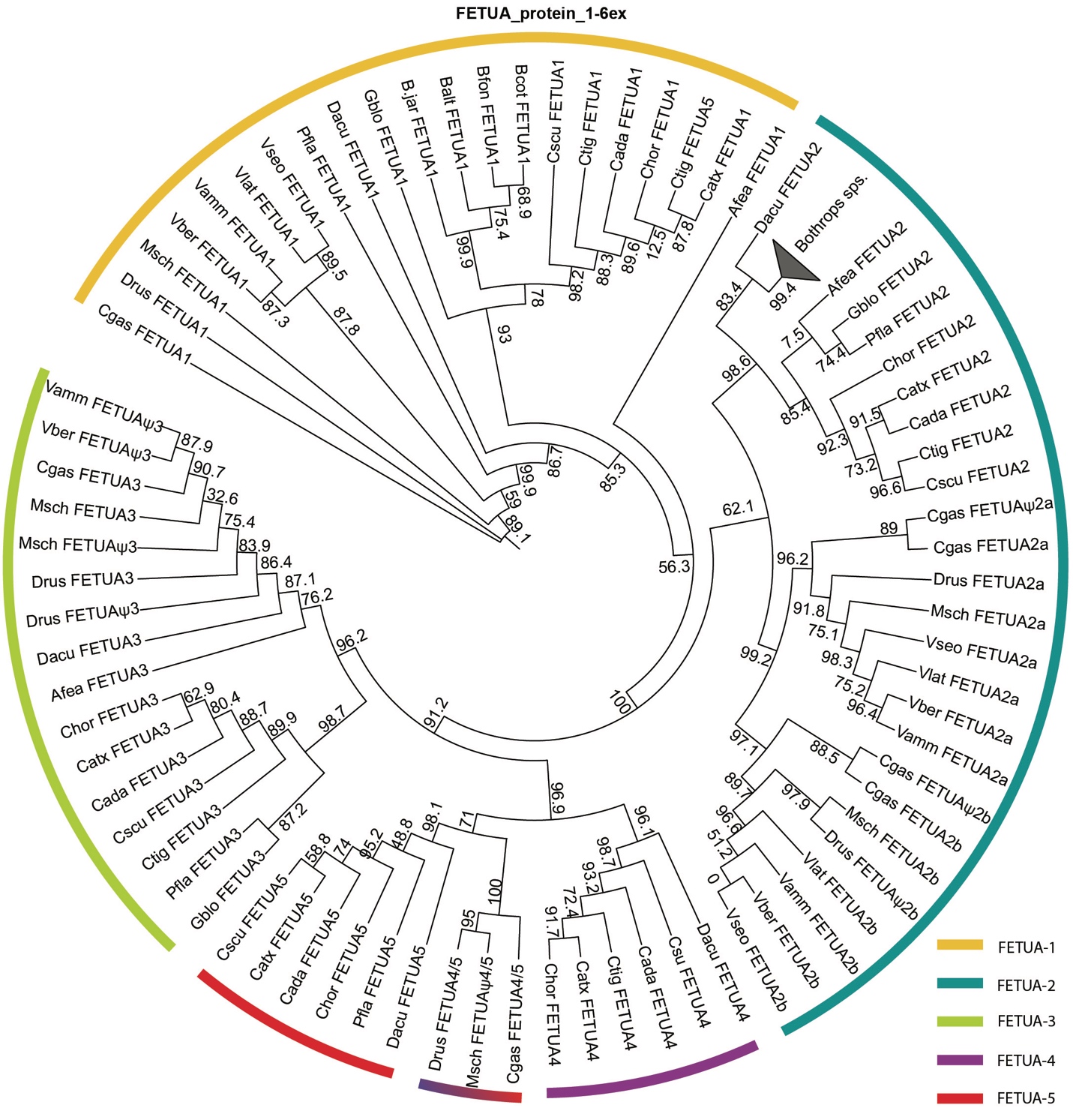
Fig. S3. All FETUA proteins belong to one of five paralog groups. Phylogenetic analysis of FETUA protein sequences from vipers. The paralog groups are indicated by colors. Exon 7 sequences were excluded due to the variable presence/absence of deletions. The newly reported Viperinae FETUA proteins include up to two members of the FETUA-2 clade, FETUA-3, and a member of the FETUA-4/FETUA-5 clade that we refer to as FETUA-4/5. Sequences used are listed in Data S1.

**A**

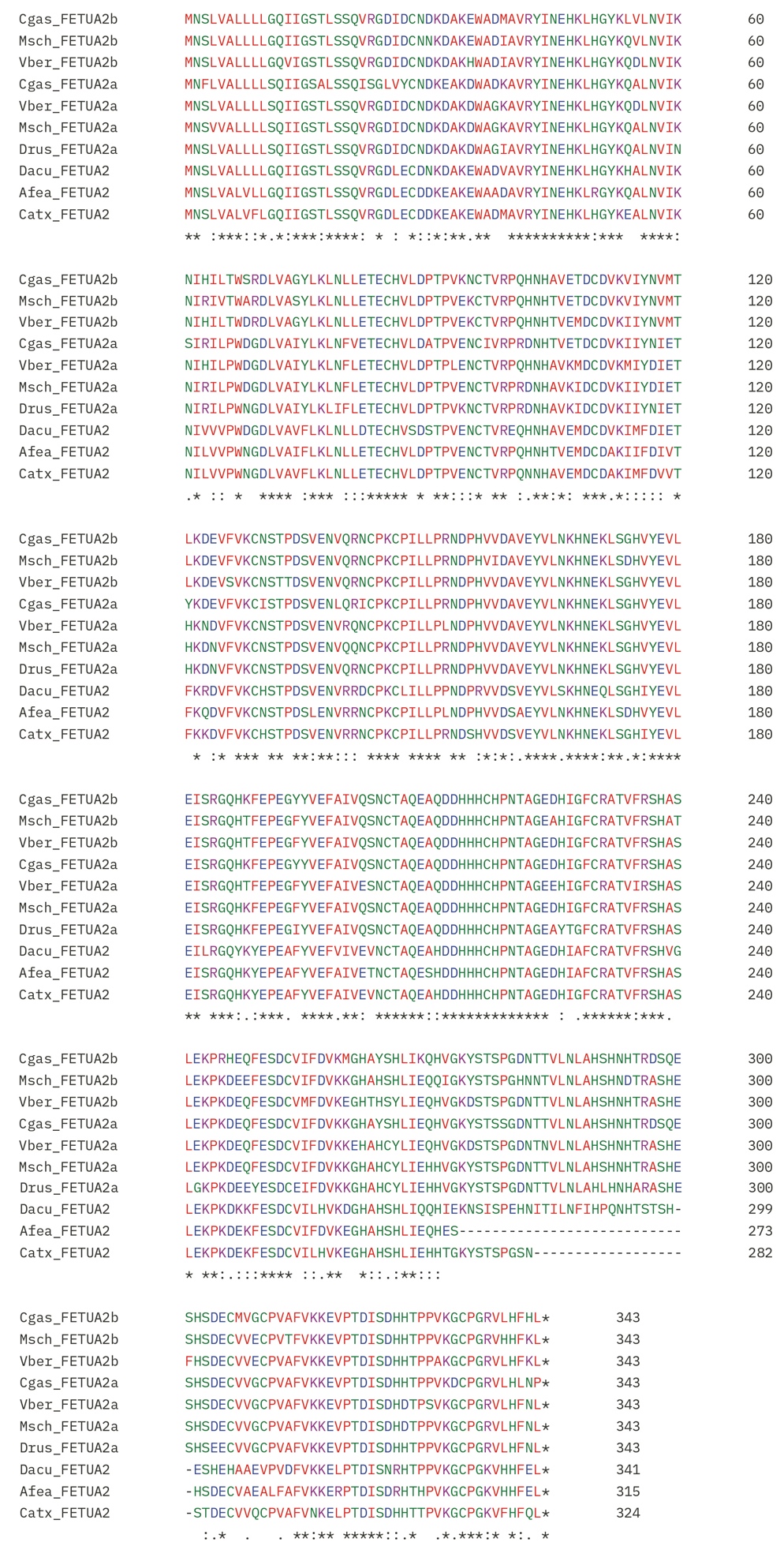

B
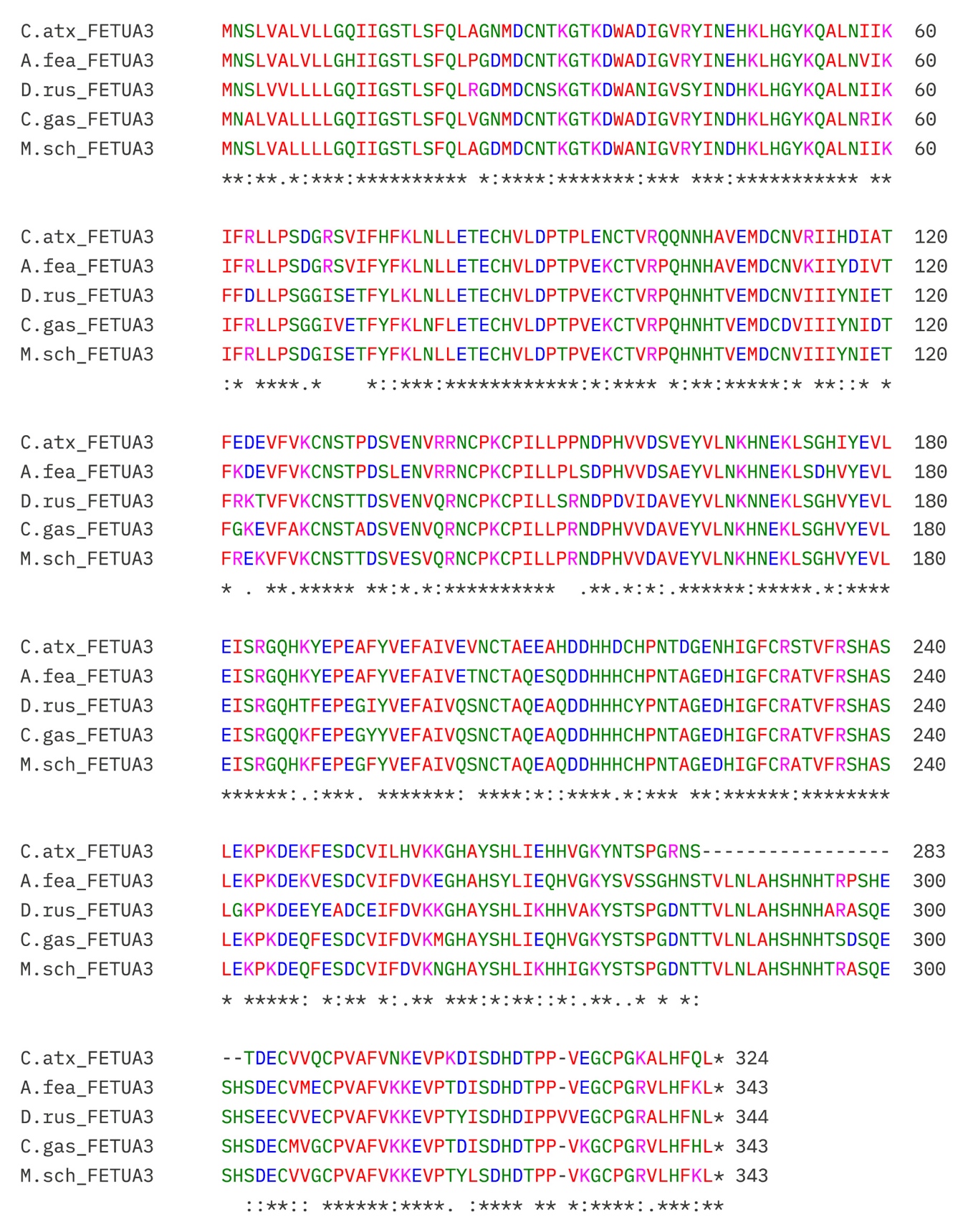

C

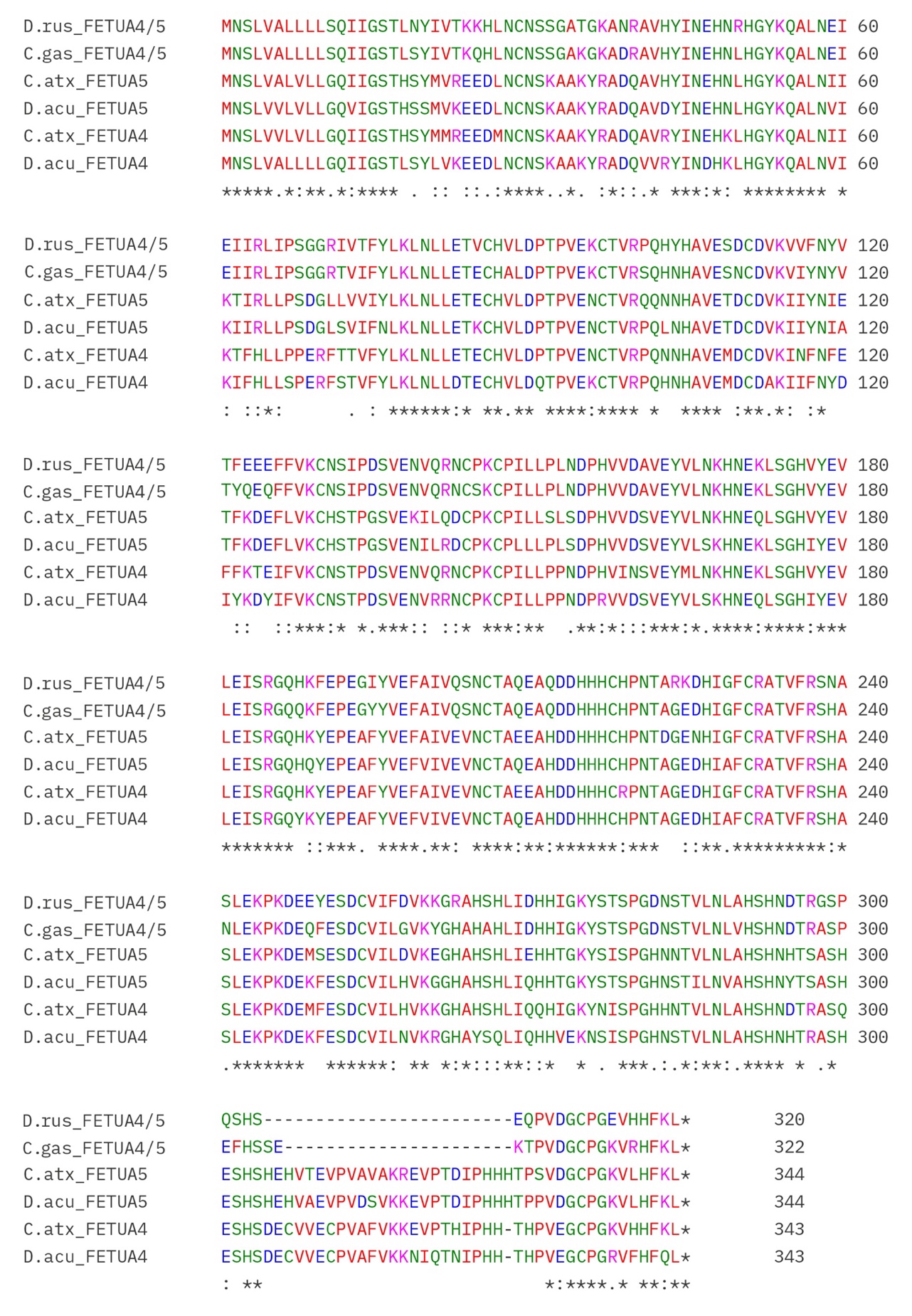

Fig. S4. Alignments of viper FETUA proteins reveal strong sequence conservation throughout each protein clade.

A. Selected proteins from the FETUA-2 clade.

The viper FETUA-2 proteins exhibit 74-77% identity to *C. atrox* FETUA-2.

B. Selected proteins from the FETUA-3 clade.

The viper FETUA-3 proteins exhibit 73-78% identity to *C. atrox* FETUA-3.

C. Selected proteins from the FETUA-4 and FETUA-5 clade.

The viper FETUA-4/5 proteins exhibit 68-70% identity to *C. atrox* FETUA-5.

Species abbreviations are M. sch, *Macrovipera schweizeri*; C. gas, *Cerastes gasperttii*; V. ber. *Vipera berus*; D. Rus, *Daboia russelli*; D. acu, *Deinagkistrodon acutus*; A. fea, *Azemiops feae;* C. atx, *Crotalus atrox*.

Symbols: Conserved amino acids are indicated by asterisks (*), sites with conservative substitutions are denoted by a colon (:), and sites with semi-conservative substitutions are denoted with a period (.)

**Dataset S1 FETUA protein sequences used in this study.**

>D.acu_FETUA1

MNSLVALLLLGQIIGCTFSHHLTSQVDCNGEDAEKWGDMAVHYINEHNLHGYKQALNVIKEIHVLPRRPHGEIVFIEIEVLETQCHVLDETPVENCTVRPQHYHAVGGDCDVKIIHHEGVDKVVGAKCHSDPDSVEDVRRNCPNCPILLPLSDPRVVDCVEYVLSKHNEKLSGHIYEVLEISRGQHQYEPEAFYVEFVIVEVNCTAQEAHDDHHHCHPNTAGEDHIAFCRATVFRSLASLEKPKDEKFESDCVILNVKEGHAHSHLIQHHVGKYSISPGHNNTVLNLAHSHNHTSASHESHSHEHVAEVPVAVAKREVPTDIPHHHMHSHPVKLCPGKVHHFKL*

>D.acu_FETUA2

MNSLVALLLLGQIIGSTLSSQVRGDLECDNKDAKEWADVAVRYINEHKLHGYKHALNVIKNIVVVPWDGDLVAVFLKLNLLDTECHVSDSTPVENCTVREQHNHAVEMDCDVKIMFDIETFKRDVFVKCHSTPDSVENVRRDCPKCLILLPPNDPRVVDSVEYVLSKHNEQLSGHIYEVLEILRGQYKYEPEAFYVEFVIVEVNCTAQEAHDDHHHCHPNTAGEDHIAFCRATVFRSHVGLEKPKDKKFESDCVILHVKDGHAHSHLIQQHIEKNSISPEHNITILNFIHPQNHTSTSHESHEHAAEVPVDFVKKELPTDISNRHTPPVKGCPGKVHHFEL*

>D.acu_FETUA3 (partial)

MNSLVALLLLGRIIGSTLSFQLAGDMDCMTKGTKDWADNVVRYINDNKLHGYKQALNVIKIFHLPPSDALSVKFYFELNFLDTECHVLDQTPVENCTVRPQNNHAVEMDCDAKIIFNYDIYKDYIFVKCNSTP

>D.acu_FETUA4

MNSLVALLLLGQIIGSTLSYLVKEEDLNCNSKAAKYRADQVVRYINDHKLHGYKQALNVIKIFHLLSPERFSTVFYLKLNLLDTECHVLDQTPVEKCTVRPQHNHAVEMDCDAKIIFNYDIYKDYIFVKCNSTPDSVENVRRNCPKCPILLPPNDPRVVDSVEYVLSKHNEQLSGHIYEVLEISRGQYKYEPEAFYVEFVIVEVNCTAQEAHDDHHHCHPNTAGEDHIAFCRATVFRSHASLEKPKDEKFESDCVILNVKRGHAYSQLIQHHVEKNSISPGHNSTVLNLAHSHNHTRASHESHSDECVVECPVAFVKKNIQTNIPHHTHPVEGCPGRVFHFQL*

>D.acu_FETUA5

MNSLVVLVLLGQVIGSTHSSMVKEEDLNCNSKAAKYRADQAVDYINEHNLHGYKQALNVIKIIRLLPSDGLSVIFNLKLNLLETKCHVLDPTPVENCTVRPQLNHAVETDCDVKIIYNIATFKDEFLVKCHSTPGSVENILRDCPKCPLLLPLSDPHVVDSVEYVLSKHNEKLSGHIYEVLEISRGQHQYEPEAFYVEFVIVEVNCTAQEAHDDHHHCHPNTAGEDHIAFCRATVFRSHASLEKPKDEKFESDCVILHVKGGHAHSHLIQHHTGKYSTSPGHNSTILNVAHSHNYTSASHESHSHEHVAEVPVDSVKKEVPTDIPHHHTPPVDGCPGKVLHFKL*

>P.fla_FETUA1

MNSLVALLLLGQIIGCTFSHHLPSHVPVDCNGEDAEKWADLAVDYINEHNLHGYKYAVNVINEIFVLPRRPHGKMILLELNLLETECHVLDQTPIKNCTVRPPHYHAVEGDCDVKIIHDEDVDKVVAAKCYSNPDSVEDVQQNCPKCPILLSLTDPHVVDSVEYVLEKHNAKLSGHIYEVLEISRGQHKYEPEAYYLEFVIVEVNCTAQEAHDDHHQCHPYTAGEEHIGFCRATVFRSHASLEKPKDEKFESDCVIFDIKEGHAHSHLIEHHVGKYSTSPGYNNTVLNLVHSHNHTSASHESHSHEHVAEVPVAVAKREVPTDIPHDHTHPVKLCPGKVHHFKL*

>P.fla_FETUA2/HSF

MNSLVALVLLGQIIGSTLSSQVRGDLECDDKEAKNWADDAVRYINEHKLHGHKQALNVIKNICVVPWNGDLVAVFLELNLLETECHVLDPTPVEKCTVRQQHNHAVEMDCDAKIMFNVETFKRDVFVKCHSTPDSVENVRRNCSKCPILLPPNNPHVVDSVEYVLNKHNEKLSGHIYEVLEISRGQHKYEPEAYYLEFVIVEINCTAQEAHDDHHQCHPYTAGEDHIAFCRSTVFRSHASLEKPKDEKFESDCVILDVKDGHAHSHLIQQHIEKNSISPEHNITILNFVHPDNHTSTSHESHEHVAEVPVVFVKKELPTDISDHHTTPVKGCPGKVHHFEL*

>P.fla_FETUA3/HLP

MNSLVALVLLGQIIGSTVSFQLGPNMDCNTKGTKDWADIGVHYINEHKLHGYKQALNVIKIFRLLPSDGRSVIFHFNLNLLETECHVLDSTPVENCTVRPQHNHAVEMDCNVRIIHDITTFEDEVFVKCSSTPGSVENILRDCPKCPILLSPNDPHVVDSVEYVLNKHNEKLSGHIYEVLEISRGQHKYEPEAYYLEFVIVEINCTAQEAHDDYHQCHPYTAGEDHIAFCRSTVFRSHASLEKPKDEKFESDCVILDVKEGHSHSHLIEHHVGKYSTSPGYNSTDECVVECPVAFVNKEVPTDISDHNTPPVKGCPGRVLHFQL*

>P.fla_FETUA5

MNSLVALVLLGQIIGSTHSYMVKEEDLNCNSKGAKY*ADQAVRYINEHNLHGYKQALNVIKIFRLLPSAGFSVIIYLKLNLLETECHVLDSTPVENCTVRQQHNHAVETDCDVKIIYNIMTFKDEFLVKCHSTPGSVENILRDCPKCPILLSLSDPHVVDSVEYVLEKHNAKLPGHIYEVFEISRGQHKYEPEALYLEFVIVEVNCTAQEAHDDHHQCHPYTAGEDHIAFCRSTVFRSHASLEKPKDEKFESDCVILDVKEGHAHSHLIQQHIEKNSISPEHNITILNFVHPHNHTSTSHESHEHVADVPVAFVKKELPTDISDHHTTPVKGCPGKVHHFEL*

>G.blo_MSF

MHFLVALVLLGQIIGSTLSSQVRGDLECNDREAKEWADQAVRYINEHKLHEYKQALNVIKNIVVVPWNGDLVAVFLKLNLLETECHVLDPTPVEKCTIRPQQNHAVEMDCDAKIMFDVETFKQDVFVKCHSTPDSVEDVRRNCLKCPILLSPSDPHVVDSVEYVLNKHNEQLSGHVYEVLEISRGQHKYEPEAFYVEFAIVEVNCTAQEAHDDHHHCHPNTAGEDHIAFCKATVFRSHASLEKPKHENFESDCVILDVKEGHAHSHLIEHHIGKYSTSPGQNSTVECVAECPVAFVNKEVPTDISDRHTTPVKGCPGKILHFQL*

>G.blo_HLP-B

MNSLVALVLLGQMIGSTLSHHLQSHVDCNGEDAEKWADMAVHYINEHNLHGYKQVFNVINEIHVLPRRPRGKIIILELKLLETECHVLDPTPVENCTVRPPHYHAVEGDCDVKILHDEGVDKVIGAKCHSDPDSVEDVRRNCPKCPILLPLSDPHVVDSVEYVLNKHNEKLSGHVYEVLEISRGQHKYEPEAFYVEFAIVEVNCTAQEAHDDHHHCHPNTAGENHIGFCRATVFRSHASLEKPKDEQFESDCVIFDVKEGHAHSHLIEHHIGNYNTSPGHNNTVLNLAHSHNHTSASHESHSHEHVAEVPVAVAKREVPTNTPHDHTHPVKLCPGKVHHFKL*

>G.blo_HLP-A

MNSLVALVLLGQIIGSTLSFQLGPNMDCNTKGTKDWADIGVRYINEHKLDGYKNALNIIKIFRLLPSDGRSVIVHFKLNLLETKCHVLDPTPVENCAVRQQHNHAVEMDCNVRIIHDIATFEDEVFVKCKSTPDSVENVRRNCPKCPILLPPNDPHVVDSVEYVLNKHNEKLSGHVYEVLEISRGQHKYEPEAFYVEFAIVEVNCTAQEARDGHHQCHPYTAGEDHIAFCRATVFRSHASLEKPKDENFESDCVILDVKEGHAHSHLIQQHIEKYSTSPGHNSTDEYVVECPVAFVEKEVPTDMSDHDTPPVKGCPGRVLHFQL*

>B.jar_FETUA1

MNSLVALVLLGQIIGSTLSHHLQSHVDCNGEDAEKWAHMAVHYINEHNQHGYKCALNVINEIRLLPRRPHGTIVFLKLKVLETECHVLDPTPTENCTVRPQHYHAVGGDCDVKIIHEEGGDKVIGAKCYSDPDSVEDVRRNCPKCPILLNLNDPQVVDSVEYVLNKHNEKVSGHVYEVLEISRGQHKNEPEAYYVEFAIVEVNCTAQEAHDDHHQCHPNTAGENHIGFCRATVFRSHASLEKPKDEQFESDCVIFDVKDGHAHSHLIEHHVGKYSTSPGHNNTVLNLVHSHNHTSASHESHSHEHVTEVPVAVAKREVPKDVPHDHTHPVKLCPGKVHHFEL*

>B.jar_FETUA2a/BJ46a

MNSLVALVLLGQIIGSTLSSQVRGDLECDEKDAKEWTDTGVRYINEHKLHGYKYALNVIKNIVVVPWDGDWVAVFLKLNLLETECHVLDPTPVKNCTVRPQHNHAVEMDCDVKIMFNVDTFKEDVFAKCHSTPDSVENVRRNCPKCPILLPSNNPQVVDSVEYVLNKHNEKLSDHVYEVLEISRGQHKYEPEAYYVEFAIVEVNCTAQELHDDHHHCHPNTAGEDHIAFCRATVFRSHASLEKPKDEQFESDCVILHVKEGHAHSHLIQQHVEKDSISPEHNNTALNFVHPHNDTSTSHESHEHLAEVPVAFVKKELPKDISDRHTTPVKGCPGKVHHFEL*

>B.jar_FETUA3

MNSMVALVLLGQIIGSTLSSQVRGDLPCDDEDSKWWADVGVRYINEHKLHGYKYALSVIKNIVVVPWDGDWVAVFLKLNLLETECHVLDPTPVKNCTVRTQHNHAVEMDCDVKIMFNVETFKEDVFAKCHSTPDSVENVRRNCPKCPILLPSNNPQVVDSVEYVLNKHNEQLSDHVYEVLEISRGQHKYEPEAYYVEFAIVEVNCTAQEAHDDHHQCHPNTAGEDHIGFCRATVFRSHASLEKPKDEQFESDCVILHVKKGHAHSHLIQQHVEKDSISPEHNNTALNFVHPHNDTRASHESHEHLAKVPVAFVKKELPKDISDRHTTPVVGCPGLRAVGQPY*

>B.jar_FETUA2b

MNSLVALVLLGQIIGSTHSYLVKEEDLNCNSKAAKYRADQAVHYINEHKLHGYKYALNVIKIFHLLPPEEFSTVFYLKLNLLETECHVFDPTPVENCNVRPQHNHAVEMDCDVKIIFNFQFFKTEVFVKCNSTPDSVENVRRNSPKCSILLPPNDPQVIDSVEYVLNKHNEQLSGHVYKVLEISRGQHKYEPEAYYVEFAIMEINCTAQEAHDDHHQCHPNTAGEDHIGFCRATVFRSHASLEKPKDEQFESDCVILDVKEGHAHSHLIQQHVEKYSTSPGHNNTALNLVHLHNDTSTSHKSHSGKCPVVFPKKKVPTDIPHHTHPVEGCPGKVLHFEL*

>C.tig_FETUA1

MNSLVALVLLGQIIGCTFSHHLQSQVDCNGEDAEKWADMAVHYINEHNLHGYKYTTNVINEIHVLPRRPHGVIVFLELKVLETQCHVLDPTPVENCTVRPQHYHAVGGDCDVKILHDEGVDKVIGAKCHSDPDSVEDVRQNCSKCPILLSLSDPHVVDSVEYVLNKYNEKLSGHVYEVLEISRGQHKYEPEAFYVEFAIVEVNCTAEEAHDDHHHCHPNTDGENHIGFCKATVFRSHASLEKPKDEMFESDCVIFDVKEGHAHSHLIEHHVGKYSTSPGHNSTVLNLVHSHNHTSASHESHSHEHVMEVPVAVAKREVPIDVPHDHTHPVKLCPGKVHHFQL*

>C.tig_FETUA2

MNSLVALVLLGXSIGSTLSSQVRGDLECDNKEAKEWADMAVRYINEHKLHGYKQALNVIKNILVVPWDGDLVAVYLKLNLLETECHMLDPHPVENCTIRPQNNHAVKMDCNAKIMFDVVTFKQDVFVKCHSTPDSVENVRRNCPKCPILLPWNDSHVVDSVEYVLNKHNEKLSGHVYEVLEISRGQHKYEPEAFYVEFAIVEVNCTAEEAHDDHHHCHPNTAGEDHIGFCRATVFRSHASLEKPKDEKFESDCVILHVKEGYAYSHLLYKQIEKYNVPPEFRNTVLNLTHSHNHTSTSHESHEHVAEVPVAFVKKELPTDISDHDTPPVEGCPGKALHFQL*

>C.tig_FETUA3

MNSLVALVLLGQIIGSTLSFQLAPNMDCNTKGTKDWADIGVRYINEHKLHGYKQALNIIKIFRLLPSDGLSVMFHFKLNLLETECHVLDPTPVENCIVRQKNNHAVEMDCNVRIIHDIATFGDEVFVKCNSTPDSVENVRRNCPKCPILLPPNDPHVVDSVEYVLNKHNEQLSGHVYEVLEISRGQHKYEPEAFYVEFAIVEVNCTAEEAHDDHHHCHPNTAGEDHIGFCRATVFRSHASLEKPKDEKFESDCVILHVKKGHAYSHLIEHHVGKYNTSPGRNSTDECVVECPVAFVNKEVPKDISDHDTPPVEGCPGKALHFQL*

>C.tig_FETUA4

MNSLVVLVLLGQIIGSTHSYMVREEHLNCNSKAAKYRADQAVRYINEHKLHGYKQALNIIKTFHLLPPERFTTVFYLKLNLLETECHVLDPTPVENCTVRPQNNHAVEMDCDVKINFNFEFFKTEIFVKCNSTPNSVENVRRNCPECPILLPPNDPHVIDSVEYVLNKHNEKLSGHVYEVLEISRGQHKYEPEAFYVEFAIVEVNCTAEEAHDDHHHCHPNTAGEDHIGFCRATVFRSHASLEKPKDEKFESDCVILHVKEGHATSHLIEHHTGKYSTSPGRNSTDECVVECPVAFVKKEVPTHIPHHTHPVEGCPGKVFHFQL*

>C.tig_FETUA5

MNSLVVLVLLGQIIGSTLSYMVREEDLNCNSKAAKYRADQAVHYINEHNLHGYKQALNIIKTIRLLPSDGLSVIIYLKLNLLETECHVLDPTPVENCTVRQQNNHAVETDCDVKIIYNIETFKDEFLVKCHSTPGSVENILQDCPKCPILMPLSEPHVTDSVEYVLNKHNEQLSGHVYEVLEISRGQHKYEPEAFYVEFAIVEVNCTAQEAHDDHHHCHPNTDGENHIGFCRATVFRSYASLEKPKVEMSESDCVILDVKEGHAHSHLIEHHVGKYSISPGHNNTVLNLAHSHNHTSASHESHSHEHVTEVPVAVAKREVPTDIPHHHTPSVDGCPGKVLHFKL*

>C.hor_FETUA5

MNSLVALVLLGQIIGSTHSYMVREEDLNCNSKAAKYRADQAVHYINEHNLHGYKQALNIIKTIRLLPSDGLSVIMYLKLNLLETECHVLDPTPVENCTVRQQHNHAVETDCDVKIIYNIETFKDEFLVKCHSTPGSVEKILEDCPKCPLLLSLSDPHVVDSVEYVLNKHNEKLSGHVYEVLEISRGQHKYEPEAFYVEFAIVEVNCTAQEAHDDHHHCHPNTAGENHIAFCRATVFRSYASLEKPKDEKFESDCVILHVKEGHTHSHLIEHHIGKYSTSPGHNSTVLNLAHSHNHTSASHESHSHEHVTEVPVAVAKREDPTDIPHHHTPSVDGCPGKVLHFKL*

>C.hor_FETUA4

MNSLVVLVLLCQIIGSTYSYMVREEDLNCNSKAAKYRADQAVRYINEHKLHGYKQALNIIKIFHLLPPERFMTVFYLKLNLLETECHVLDPTPVENCTVRPQNNHAVEMDCDVKINFNFEFFKTEIFVKCNSTPDSVENVQRNCPECPILLPPNDPHVINSVEYVLNKHNEKLSGHVYEVLEISRGQHKYEPEAFYVEFAIVEVNCTAEEAHNDHHHCHPNTAGKDHIGFCRATVFRSHASLEKPKDEMFESDCVIFDVKKGHAHSHLIQQHIGKYNISPGHNNAVLNLAHSHNDTRASQESHSDECVVECPVAFVKKEVPTHIPHHTHPVEGCPGKVFHFQL*

>C.hor_FETUA3

MNSLVALMLLGQIIGSTLSFQLAANMDCNTKGTKDWADIGVRYINEHKLHGYKQDLNIIKIFRLLPSDGRSVIFHFKLNLLETECHVLDPTPVENCTVRQQNNHAVEMDCNVRIIHDIATFEDEVFVKCNSTPDSVENVRRNCPKCPILLPPNDPHVVDSVEYVLNKHNEKLSGHIYEVLEISRGQHKYEPEAFYVEFAIVEVNCTAEEAHDDHHHCHPNTDGENHIGFCRSTVFRSHASLEKPKDEKFESDCVILHVKEGHAHSHLIEHHVGKYSTSPGRNSTDECVVECPVAFVNKEVPKDISDHDTPPVEGCPGKVLHFQL*

>C.hor_FETUA2

MNSLVALVLLGQIIGSTLSSQVRGDLECDDKEAKEWADIAVRYINEHKLHGYKQALNVIKNILVVPWNGDLVAVFLKLNLLETECHVLDPTPVENCTVRPQNNHAVEMDCDAKIMFDVVTFKTDVFVKCHSTPDSVENVRRNCPKCPILLPRNDSHVVDSVEYVLNKHNEKLSGHVYEVLEISRGQHKSEPEAFYVEFAIVEVNCTAQEAHDDHHHCHPNTTGEDHIGFCRATVFRSHASLEKPKDEKFESDCVIFDVKEGHAHSHLIEHHVGKYSTSPGRNSTDECVVECPVAFVNKELPTDISDHHTTPVKGCPGKVLHFQL*

>C.hor_FETUA1

MNSLVALVLLGQIIGCTFSHHLQSQVDCNGEDAEKWADMAVHYINEHNLHGYKYTNNVINEIHVLPRRPRGEIVFLELKVLETQCHVLDPTPVENCTVRPLHYHAVGGDCDVKIIHDEGVDKVIGAKCHSDPDSVEEVRRNCPECPILLSLSDPHVINCVEYVLNKYNEKLSGHVYEVLEISRGQHKYEPEAFYVEFAIVEVNCTAEEAHDDHHHCHPNTDGENHIGFCRATVFRSHASLEKPKDEMFESDCVIFDVKEGHAHSHLIEHHVGKYSTSPGHNNTVLNLIHSHNHTSASHESHSHEHMTEVPVAVAKREVPTDVPHDHIHPVKLCPGKVHHFKL*

>C.scu_FETUA5

MNSLVALVLLGQIIGSTHSYMVREEDLNCNSKAAKYRADQAVHYINEHNLHGYKQALNIIKTIRLLPSDGLSVIIYLKLNLLETECHVLDPTPVENCTVRQQNNHAVETDCDVKIIYNIETFKDEFLVKCHSTPGSVEKILQDCPKCPILLSLSDPHVVDSVEYVLNKHNEQLSGHVYEVLEISRGQHKYEPEAFYVEFAIVEVNCTAQEAHDDHHHCHPNTAGEDHIAFCRATVFRSYASLEKPKDEMSESDCVILDVKEGHAHSHLIEHHTGKYSTSPGHNSTVLNLAHSHNHTSASHESHSHEHVTEVPVAVAKREVPTDIPHHHTPSVDGCPGKVLHFKL*

>C.scu_FETUA4

MNSLVALVLLGQIIGSTLSYMVREEDLNCNSKAAKYRADQAVRYINEHKLHGYKQALNIIKTFHLLPPERFTTVFYLKLNLLETECHVLDPTPVENCTVRPQYNHAVEMDCDVKINFNFEFFKTEIFVKCNSTPDSVENVR*NCPECPILLPPNDPHVIDSVEYVLNKHNEKLSGHIYEVLEISRGQHKYEPEAFYVEFAIVEVNCTAEEAHDDHHHCHPNTAGEDHIGFCRATVFRSHASLEKPKDEKFESDCVILHVKKGHAHSHLIQQHIGKYNISPGHHNTVLNLAHSHNDTRASQESHSDECVVECPVAFVKKEVPTHIPHHTHPVEGCPGKVHHFKL*

>C.scu_FETUA3

DGLSVIFHFKLNLLETECHVLDPTPLENCTVRQQNNHAVEMDCNVRIIHDIATFEDEVFVKCNSTPDSVENVRRNCPKCPILLPPNDPHVVDSVEYVLNKHNEKLSGHIYEVLEISRGQHKYEPEAFYVEFAIVEVNCTAEEAHDDHHHCHPNTDGENHIGFCRATVFRSHASLEKPKDEKFESDCVILHVKKGHAYSHLIEHHVGKYNTSPGRNSTDECVVECPVAFVNKEVPKDISDHDTPPVEGCPGKALHFQL*

>C.scu_FETUA2

MNSLVALVLLGQIIGSTLSSQVRGDLECDDKEAKEWADMAVRYINEHKLHGYKQALNVIKNILVVPWDGDLVAVYLKLNLLETECHVLDPTPVENCTIRPQNNHAVKMDCNAKIMFDVVTFKQDVFVKCHSTPDSVENVRRNCPECPILLPRNDSHVVDSVEYVLNKHNEKLSGHVYEVLEISRGQHKYEPEAFYVEFAIVEVNCTAQEAHDDHHHCHPNTAGENHIGFCRATVFRSHASLEKPKDEKFESDCVILHVKEGHAHSHLIEHHVGKYNTSPGRNSTDECVVESSLLTKATGRPVAFVNKEVPADISDHHTTPVKGCPGKVLHFQL*

>C.scu_FETUA1

MNSLVALVLLGQIIGCTFSHHLQSQVDCNGEDAEKWADMAVHYINEHNLHGYKYTTNVINEIHVLPRRPHGEIIFLELKVLETQCHVLDPTPVENCTVRPLHYHAVGGDCDVKILHDEGVDKVIGAKCHSDPDSVEDVRRNCPNCPILLSLSDPHVVDSVEYVLNKHNEKLSGHVYEVLEISRGQHKYEPEAFYVEFAIVEVNCTAEEAHDDHHHCHPNTAGENHIGFCRATVFRSHASLEKPKDEKFESDCVIFDVKEGHAHSHLIEHHTGKYSTSPGHNNTVLNLVHSHNHTSASHESHSHEHMTEVPVAVAKREVPTDVPHDHTHPVKLCPGKVHHFKL*

>C.atx_FETUA5

MNSLVALVLLGQIIGSTHSYMVREEDLNCNSKAAKYRADQAVHYINEHNLHGYKQALNIIKTIRLLPSDGLLVVIYLKLNLLETECHVLDPTPVENCTVRQQNNHAVETDCDVKIIYNIETFKDEFLVKCHSTPGSVEKILQDCPKCPILLSLSDPHVVDSVEYVLNKHNEQLSGHVYEVLEISRGQHKYEPEAFYVEFAIVEVNCTAEEAHDDHHHCHPNTDGENHIGFCRATVFRSHASLEKPKDEMSESDCVILDVKEGHAHSHLIEHHTGKYSISPGHNNTVLNLAHSHNHTSASHESHSHEHVTEVPVAVAKREVPTDIPHHHTPSVDGCPGKVLHFKL*

>C.atx_FETUA4

MNSLVVLVLLGQIIGSTHSYMMREEDMNCNSKAAKYRADQAVRYINEHKLHGYKQALNIIKTFHLLPPERFTTVFYLKLNLLETECHVLDPTPVENCTVRPQNNHAVEMDCDVKINFNFEFFKTEIFVKCNSTPDSVENVQRNCPKCPILLPPNDPHVINSVEYMLNKHNEKLSGHVYEVLEISRGQHKYEPEAFYVEFAIVEVNCTAEEAHDDHHHCRPNTAGEDHIGFCRATVFRSHASLEKPKDEMFESDCVILHVQKGHAHSHLIQQHIGKYNISPGHHNTVLNLAHSHNDTRASQESHSDECVVECPVAFVKKEVPTHIPHHTHPVEGCPGKVHHFKL*

>C.atx_FETUA3

MNSLVALVLLGQIIGSTLSFQLAGNMDCNTKGTKDWADIGVRYINEHKLHGYKQALNIIKIFRLLPSDGRSVIFHFKLNLLETECHVLDPTPLENCTVRQQNNHAVEMDCNVRIIHDIATFEDEVFVKCNSTPDSVENVRRNCPKCPILLPPNDPHVVDSVEYVLNKHNEKLSGHIYEVLEISRGQHKYEPEAFYVEFAIVEVNCTAEEAHDDHHDCHPNTDGENHIGFCRSTVFRSHASLEKPKDEKFESDCVILHVKKGHAYSHLIEHHVGKYNTSPGRNSTDECVVQCPVAFVNKEVPKDISDHDTPPVEGCPGKALHFQL*

>C.atx_FETUA2

MNSLVALVFLGQIIGSTLSSQVRGDLECDDKEAKEWADMAVRYINEHKLHGYKEALNVIKNILVVPWDGDLVAVFLKLNLLETECHVLDPTPVENCTVRPQNNQAVEMDCDAKIMFDVVTFKKDVFVKCHSTPDSVENVRRNCPKCPILLPRNDSHVVDSVEYVLNKHNEKLSGHIYEVLEISRGQHKYEPEAFYVEFAIVEVNCTAQEAHDDHHHCHPNTAGEDHIGFCRATVFRSHASLEKPKDEKFESDCVILHVQEGHAHSHLIEHHTGKYSTSPGSNSTDECVVQCPVAFVNKELPTDISDHHTTPVKGCPGKVFHFQL*

>C.atx_FETUA1

MNSLVALVLLGQIIGCTFSHHLQSQIDCNGEDAEKWADMAVHYINEHNLHGYKYTTNVINEIHVLPWRPHGEIIFLELKVLETQCHVLDPTPVENCTVRPLHYHAVGGDCDVKIIHDEGVDKVIGAKCHSDPDSVENVRQNCPECPILLSLSDPHVVDCVEYVLNKYNEKLSGHIYEVLEISRGQHKYEPEAFYVEFAIVEVNCTAEEAHDDHHHCHPNTDGENHIGFCRATVFRSHASLEKPKDEMFESDCVIFDVKEGHAHSHLIEHHTGKYSTSPGHNNTVLNLIHSHNHTSASHESHSDEHMTEVPVAVAKREVPTDVPHDHTHPVKLCPGKVHHFKL*

>C.ada_FETUA5

MNSLVALVLLGQIIGSIHSYMVREEDLNCNSKAAKYRADQAVQYINEHNLHGYKQALNIIKTIRLLPSDGLLVVIYLKLNLLETECHVLDPTPVENCTVRPQNNHAVETDCDVKIIYNIETFKDEFLVKCHSTPGSVEKILQDCPKCPILLPPNDPHVVDVEYVLNKHNEKLSGHVYEVLEISRGQHKYEPEAFYVEFAIVEVNCTAQEAHDDHHHCHPNIAGEDHIGFCRATVFRSYASLEKPKDEKFESDCVILDVKEGHAHSHLIQQHIGKYSTSPGHNNTVLNLAHSHNHTSASHESHSHEHVTEVPVAVAKREDPTDIPHHHTPSVDGCPGKVLHFKL*

>C.ada_FETUA4

MNSLVVLVLLGQIIGSTHSYMVREEDMNCNSKAAKYRADQAVRYINEHKLHGYKQALNIIKTFHLLPPERFTTVFYLKLNLLETECHVLDPTPVENCTVRPQNNHAVEMDCDVKINFNFEFFKTEIFVKCNSTPDSVENVRRNCPECPILLPPNDPHVIDSVEYVLNKHNEKLSGHIYEVLEISRGQHKYEPEAFYVEFAIVEVNCTAEEAHDDHHHCHPNTDGENHIGFCRATVFRSHASLEKPKDEKFESDCVILHVKEGHAHSHLIQQHIGKYNISPGHNNTVLNLAHSHNDTRASQESHSDECVVECPVVFVKKEVPTHIPHHTHPVEGCPGKVFHFQL*

>C.ada_FETUA3

MNSLVALMLLGQIIGSTLSFQLAGNMDCNTKGTKDWADIGVRYINEHKLHGYKQALNIIKIFRLLPSGGLSVIFHFKLNLLETECHVLDPTPVENCTVRPQNNHAVEMDCNVRIIHDIATFEDEVFVKCNSTPDSLENVRRNCPKCPILLPPNDPHVVDSVEYVLNKHNEKLSGHIYEVLEISRGQHKYEPEAFYVEFAIVEVNCTAEEAHDDHHHCHPNTDGENHIGFCRATVFRSHASLEKPKDEKFESDCVILHVKEGHAHSHLIQHHTGKYSTSPGRNSTDECVVECPVAFVNKELPTDISDHDTPPVEGCPGKVLHFQL*

>C.ada_FETUA2

MNSLVALVFLGQIIGSTLSSQVRGDLECDDKEAKEWADMAVRYINEHKLHGYKEALNVIKNILVVPWNGDLVAVFLKLNLLETECHVLDPTPVENCTVRPQNNHAVEMDCDAKIMFDVVTFKKDVFVKCLSTPDSVENVRRNCPKCPILLPRNDSHVVDSVEYVLNKHNEKLSGHVYEVLEISRGQHKYEPEAFYVEFAIVEVNCTAQEAHDDHHHCHPNTAGEDHIGFCRATVFRSHASLEKPKDEKFESDCVILHVKEGHAHSHLIEHHTGKYSTSPGRNSTDECVVQCPVAFVNKELPTDISDHHTTPVKGCPGKVFHFQL*

>C.ada_FETUA1

MNSLVALVLLGQIIGCTFSHHLQSQIDCNGEDAEKWADMAVHYINEHNLHGYKYTTNVINEIHVLPRRPHGEIIFLELKVLETQCHVLDPTPVENCTVRPLHYHAVGGDCDVKIIHDEGVDKVIGAKCHSDPDSVENVRQNCPECPILLSLSDPHVIDSVEYVLNKYNEKLSGHVYEVLEISRGQHKYEPEAFYVEFAIVEVNCTAEEAHDDHHHCHPNTDGENHIGFCRATVFRSHATLEKPKDEMFELDCVIFDVKEGHAHSHLIEHHVGKYSTSPGHNNTVLNLIHSHNHTSASHESHSHEHMMEVPVAVAKREVPTDVPHDHTHPVKLCPGKVHHFKL*

>Drus_FETUA1

MNSLVALLLLGQIIGCTFSHHLPSHGDCNGEDAKKWANLAVDYINEHTLHGYKQELNIIKDIHVLPRRPHGKIAFLKLELLETECHVLDKTPVKNCTVRPQQNHAVEGECDVKIIHDEDVDKVVAVKCHSSPDSVEDVRRNCPNCPILLRLNDPRVVDAVEYVLNKVNEKLPSYIYELLEITRGQLTFEPEGFYVEFAIVQSNCTAQEAQDDHHQCHPNTEEEDHTGFCKATLFRSHDSLGKPKDEQFQPDCELFDVKEGLAYSHLIEHRDGKYTSPGYNSTVLNLAHSHNHTSASQESHSHEHVVEVAVAVAKREIPTDIPPHHTDPVNLCPGEVYHFTL*

>Drus_FETUA4/5

MNSLVALLLLSQIIGSTLNYIVTKKHLNCNSSGATGKANRAVHYINEHNRHGYKQALNEIEIIRLIPSGGRIVTFYLKLNLLETVCHVLDPTPVEKCTVRPQHYHAVESDCDVKVVFNYVTFEEEFFVKCNSIPDSVENVQRNCPKCPILLPLNDPHVVDAVEYVLNKHNEKLSGHVYEVLEISRGQHKFEPEGIYVEFAIVQSNCTAQEAQDDHHHCHPNTARKDHIGFCRATVFRSNASLEKPKDEEYESDCVIFDVKKGRAHSHLIDHHIGKYSTSPGDNSTVLNLAHSHNDTRGSPQSHSEQPVDGCPGEVHHFKL*

>Drus_FETUA3

MNSLVVLLLLGQIIGSTLSFQLRGDMDCNSKGTKDWANIGVSYINDHKLHGYKQALNIIKFFDLLPSGGISETFYLKLNLLETECHVLDPTPVEKCTVRPQHNHTVEMDCNVIIIYNIETFRKTVFVKCNSTTDSVENVQRNCPKCPILLSRNDPDVIDAVEYVLNKNNEKLSGHVYEVLEISRGQHTFEPEGIYVEFAIVQSNCTAQEAQDDHHHCYPNTAGEDHIGFCRATVFRSHASLGKPKDEEYEADCEIFDVKKGHAYSHLIKHHVAKYSTSPGDNTTVLNLAHSHNHARASQESHSEECVVECPVAFVKKEVPTYISDHDIPPVVEGCPGRALHFNL*

>Drus_FETUA2a

MNSLVALLLLSQIIGSTLSSQVRGDIDCNDKDAKDWAGIAVRYINEHKLHGYKQALNVINNIRILPWNGDLVAIYLKLIFLETECHVLDPTPVKNCTVRPRDNHAVKIDCDVKIIYNIETHKDNVFVKCNSTPDSVENVQRNCPKCPILLPRNDPHVVDAVEYVLNKHNEKLSGHVYEVLEISRGQHKFEPEGIYVEFAIVQSNCTAQEAQDDHHHCHPNTAGEAYTGFCRATVFRSHASLGKPKDEEYESDCEIFDVKKGHAHCYLIEHHVGKYSTSPGDNTTVLNLAHLHNHARASHESHSEECVVGCPVAFVKKEVPTDISDHHTPPVKGCPGRVLHFNL*

>Cgas_FETUA1

MNSLVALLLVGQIIGSTLSHHLPSHGDCNGEDAEKWAHLAVHYINEHNLHGYKQDLNIIKDIHVLPRRPHGKIVFLKLELLETVCHVLDPTPVKNCTVRPQHYHAVEGDCDVKILHDENEDNVVAVKCHSSPDSVEDVQRNCPNCPILLRLNDPHVVDAVEYVLNKHNEKLSGHVYEVLEISRGQHKFEPEGYYVEFAIVQSNCTAQEAQDDHHHCHPNTAGEDHIGFCRATVFRSHASLEKPKDEQFESDCVIFNVKEGHAHSHLIEHHVGKYSTSPGHNNTVLNLAHSHNHTSASHESHSHEVPVAVAKREVPTNISPPHTHPVNLCPGKVHHFKV*

>Cgas_FETUA4/5

MNSLVALLLLSQIIGSTLSYIVTKQHLNCNSSGAKGKADRAVHYINEHNLHGYKQALNEIEIIRLIPSGGRTVIFYLKLNLLETECHALDPTPVEKCTVRSQHNHAVESNCDVKVIYNYVTYQEQFFVKCNSIPDSVENVQRNCSKCPILLPLNDPHVVDAVEYVLNKHNEKLSGHVYEVLEISRGQQKFEPEGYYVEFAIVQSNCTAQEAQDDHHHCHPNTAGEDHIGFCRATVFRSHANLEKPKDEQFESDCVILGVKYGHAHAHLIDHHIGKYSTSPGDNSTVLNLVHSHNDTRASPEFHSSEKTPVDGCPGKVRHFKL*

>Cgas_FETUA3

MNALVALLLLGQIIGSTLSFQLVGNMDCNTKGTKDWADIGVRYINDHKLHGYKQALNRIKIFRLLPSGGIVETFYFKLNFLETECHVLDPTPVEKCTVRPQHNHTVEMDCDVIIIYNIDTFGKEVFAKCNSTADSVENVQRNCPKCPILLPRNDPHVVDAVEYVLNKHNEKLSGHVYEVLEISRGQQKFEPEGYYVEFAIVQSNCTAQEAQDDHHHCHPNTAGEDHIGFCRATVFRSHASLEKPKDEQFESDCVIFDVKMGHAYSHLIEQHVGKYSTSPGDNTTVLNLAHSHNHTSDSQESHSDECMVGCPVAFVKKEVPTDISDHDTPPVKGCPGRVLHFHL*

>Cgas_FETUA2a

MNFLVALLLLSQIIGSALSSQISGLVYCNDKEAKDWADKAVRYINEHKLHGYKQALNVIKSIRILPWDGDLVAIYLKLNFVETECHVLDATPVENCIVRPRDNHTVETDCDVKIIYNIETYKDEVFVKCISTPDSVENLQRICPKCPILLPRNDPHVVDAVEYVLNKHNEKLSGHVYEVLEISRGQHKFEPEGYYVEFAIVQSNCTAQEAQDDHHHCHPNTAGEDHIGFCRATVFRSHASLEKPKDEQFESDCVIFDVKKGHAYSHLIEQHVGKYSTSSGDNTTVLNLAHSHNHTRDSQESHSDECVVGCPVAFVKKEVPTDISDHHTPPVKDCPGRVLHLNP*

>Cgas_FETUA2b

MNSLVALLLLGQIIGSTLSSQVRGDIDCNDKDAKEWADMAVRYINEHKLHGYKLVLNVIKNIHILTWSRDLVAGYLKLNLLETECHVLDPTPVKNCTVRPQHNHAVETDCDVKVIYNVMTLKDEVFVKCNSTPDSVENVQRNCPKCPILLPRNDPHVVDAVEYVLNKHNEKLSGHVYEVLEISRGQHKFEPEGYYVEFAIVQSNCTAQEAQDDHHHCHPNTAGEDHIGFCRATVFRSHASLEKPRHEQFESDCVIFDVKMGHAYSHLIKQHVGKYSTSPGDNTTVLNLAHSHNHTRDSQESHSDECMVGCPVAFVKKEVPTDISDHHTPPVKGCPGRVLHFHL*

>Msch_FETUA1

MNSLVALLLLGQIIGCTFSHHLPSHGDCNGEDAKKWAHLAVHYINEHNLHGYKQDLNIIKDIHVLPRRPHGKIVFLELKLLETVCHVLDPTPVENCTVRPQHYHVAVEGDCDVKIIHDEDVDKVVAAKCHSSPDSVEDVRRNCPNCPILLRLNDPHVVDTVEYVLNKHNEKLSGHVYEVLEISRGQHTFEPEGFYVEFAIVQSNCTAQEAQDDHHHCHPNTAGEDHIGFCRATVIRSHASLEKPKDEQFESDCVIFDVKEGHSYSHLIEHHVGKYSTSPGHNNTVLNLAHSHNHTSASHESHSHEHVVEVPVAVAKREIPTDIPPHHTHPVNLCPGKVHHFKV*

>Msch_FETUA3

MNSLVALLLLGQIIGSTLSFQLAGDMDCNTKGTKDWANIGVRYINDHKLHGYKQALNIIKIFRLLPSDGISETFYFKLNLLETECHVLDPTPVEKCTVRPQHNHTVEMDCNVIIIYNIETFREKVFVKCNSTTDSVESVQRNCPKCPILLPRNDPHVVDAVEYVLNKHNEKLSGHVYEVLEISRGQHKFEPEGFYVEFAIVQSNCTAQEAQDDHHHCHPNTAGEDHIGFCRATVFRSHASLEKPKDEQFESDCVIFDVKNGHAYSHLIKHHIGKYSTSPGDNTTVLNLAHSHNHTRASQESHSDECVVGCPVAFVKKEVPTYLSDHDTPPVKGCPGRVLHFKL*

>Msch_FETUA2a

MNSVVALLLLSQIIGSTLSSQVRGDIDCNDKDAKDWAGKAVRYINEHKLHGYKQALNVIKNIRILPWDGDLVAIYLKLNFLETECHVLDPTPVENCTVRPRDNHAVKIDCDVKIIYDIETHKDNVFVKCNSTPDSVENVQQNCPKCPILLPRNDPHVVDAVEYVLNKHNEKLSGHVYEVLEISRGQHKFEPEGFYVEFAIVQSNCTAQEAQDDHHHCHPNTAGEDHIGFCRATVFRSHASLEKPKDEQFESDCVIFDVKKGHAHCYLIEHHVGKYSTSPGDNTTVLNLAHSHNHTRASHESHSDECVVGCPVAFVKKEVPTDISDHDTPPVKGCPGRVLHFNL*

>Msch_FETUA2b

MNSLVALLLLGQIIGSTLSSQVRGDIDCNNKDAKEWADIAVRYINEHKLHGYKQVLNVIKNIRIVTWARDLVASYLKLNLLETECHVLDPTPVEKCTVRPQHNHTVETDCDVKVIYNVMTLKDEVFVKCNSTPDSVENVQRNCPKCPILLPRNDPHVIDAVEYVLNKHNEKLSDHVYEVLEISRGQHTFEPEGFYVEFAIVQSNCTAQEAQDDHHHCHPNTAGEAHIGFCRATVFRSHATLEKPKDEEFESDCVIFDVKKGHAHSHLIEQQIGKYSTSPGHNNTVLNLAHSHNDTRASHESHSDECVVECPVTFVKKEVPTDISDHHTPPVKGCPGRVHHFKL*

>Vseo_FETUA1

MNSLVALLLLGQIIGCTFSHHLPSHGDCNGEDAKKWAHLAVHYINKHTLHGYKQDLNIIKDIHVLPRRPHGKIVFLELELLETVCHVLDPTPVENCTVRPQHYHAVEGDCDVKIIHDEDVDNVVAAKCHSSPDSVEDVRQNCPKCPILLPLNDPHVVDAVEYVLNKHNEKLSGHVYEVLEISRGQHTFEPEGFYVEFAIVESNCTDQEAQDDHHHCHPNTAGEDHIGFCRANVIRSHASLEKPKDEQFESDCVIFDVKEGHSYSHLIEHHVGKYSTSPGHNNTVLNLAHSHNHTSASHESHSHEHVVEVPVAVAKREIPTNIPPHHTHPVNLCPGKVHHFKV*

>Vseo_FETUA2a

MNSLVALLLLSQIIGSTLSSQVRGDIDCNDKDAKDWAGKAVRYINEHKLHGYKQALNVIKNIHILPWDGDLVAIYLKLNFLETECHVLDPTPVENCTVRPQHNHAVKIDCDVKIIYDIETHKDDVFVKCNSTPDSVENVQQNCPKCPILLPLNDPHVVDAVEYVLNKHNEKLSGHVYEVLEISRGQHKFEPEGFYVEFAIVESNCTAQEAQDDHHHCHPNTEGEDHIGFCRATVFRSHASLEKPKDEQFESDCVIFDVKKGHAHCYLIEQHVGKDSTSPGDNTNVLNLAHSHNHTRASHESHSDECVVGCPVAFVKKEVPTDISDHDTPSVKGCPGRVLHFNL*

>Vseo_FETUA2b

MNSLVALLFLGQVIGSTLSSQVRGDIDCNDKDAKHWADIAVRYINEHKLHGYKQDLNVIKNIHILTWDRDLVAGYLKLNLLETECHVLDPTPVEKCTVRPQHNHTVEMDCDVKIIYNVMTLKDEVSVKCNSTTDSVENVQRNCPKCPILLPRNDPHVVDAVEYVLNKHNEKLSGHVYEVLEISRGQHTFEPEGFYVEFAIVQSNCTAQEAQDDHHHCHPNTAGEDHIGFCRATVFRSHASLEKPKDEQFESDCVMFDVKEGHTHSYLIEQHIGKDSTSPGDNTTVLNLAHSHNHTRTSHEFHSDECVVECPVAFIKKEVPTDISDHHTPPVKGCPGRVLHFKL*

>Vamm_FETUA1

MNSLIALLLLGQIIGCTFSHHLPSHGDCNGEDAKKWAHLAVHYINEHTLHGYKQDLNIIKDIHVLPRRPRGKIIFLELELLETVCHVLDPTPVENCTVRPQHYHAVEGDCDVKIIHDEDVDKVVAAKCHSSPDSVEDVRRNCPKCPILLPLNDPHVVDAIKYVLNKHNEKLSGHVYEVLEISRGQYTFEPEGFYVEFAIVQSNCTAQEAQDDHHHCHPNTAGEDHIGFCRANVIRSHASLEKPKDEQFESDCVIFDVKEGHSYSHLIEHHVGKYSTSPGHNNTVLNLAHSHNHTSASHESHSHEHVVEVPVAVAKREIPTNIPPHHTHPVNLCPGKVHHFKV*

>Vamm_FETUA2a

MNFVVALLLLSQIIGSTLSSQVRGDIDCNDKDAKDWAGKAVRYINEHKLHGYKQDLNVIKNIHILPWNGDLVAIYLKLNFLETECHVLDPTPLENCTVRPRDNHAVKMDCDVKMIYDIETHKNDVFVKCNSTPDSVEHVQQNCPKCPILLPQNDPHVVDAVEYVLNKHNEKLSGHVYEVLEISRGQHTFEPEGFYVEFAIVESNCTAQEAQDDHHHCHPNTEGEDHIGFCRATVIRSHASLEKPKDEQFESDCVIFDVKKGHAHCYLIEQHVGKDSTSPGDNTTVLNLAHSHNHTRASHESHSDECVVGCPVAFVKKEVPTDISDHDTPSVKGCPGRVLHFNL*

>Vamm_FETUA2b

MNSLVALLLLGQIIGSTLSSQVRGDIDCNDKDAKHWADIAVRYINEHKLHGYKQDLNVIKNIHILTWDRDLVAGYLKLNLLETECHVLDPTPVEKCTVRPQHNHTVEMDCDVKIIYNVMTLKDEVSVKCNSTTDSVENVQRNCPKCPILLPRNDPHVVDAVEYVLNKHNEKLSGHVYEVLEISRGQHTFEPEGFYVEFAIVQSNCTAQEAQDDHHHCHPNTEGEDHIGFCRATVFRSHASLEKPKDEQFESDCVMFDVKEGHAHSYLIEQHVGKDSTSPGDNTTVLNLAHSHNHTRASHEFHSDECVVECPVAFVKKEVPTDISDHHTPPAKGCPGRVLHFKL*

>Vber_FETUA1

MNSLIALLLLGQIIGCTFSHHLPSHGDCNGEDAKKWAHLAVHYINEHTLHGYKQDLNIIKDIHVLPRRPRGKIIFLELELLETVCHVLDPTPLENCTVRPQHYHAVEGDCDVKIIHDEDVDKVVAAKCHSSPDSVEDVRQNCPKCPILFPLNDPHVVDAVEYVLNKHNEKLSGHVYEVLEISRGQYTFEPEGFYVEFAIVESNCTAQEAQDDHHHCHPNTAGEEHIGFCRANVVRSHASLEKLKDEQFESDCVIFDVKEGHSYSHLIEHHIGKYSTSPGHNNTVLNLAHSHNHTSASHESHSHEHVVEVPVAVAKREIPTNIPPHHTHPVNLCPGKVHHFKV*

>Vber_FETUA2a

MNSLVALLLLSQIIGSTLSSQVRGDIDCNDKDAKDWAGKAVRYINEHKLHGYKQDLNVIKNIHILPWDGDLVAIYLKLNFLETECHVLDPTPLENCTVRPQHNHAVKMDCDVKMIYDIETHKNDVFVKCNSTPDSVENVRQNCPKCPILLPLNDPHVVDAVEYVLNKHNEKLSGHVYEVLEISRGQHTFEPEGFYVEFAIVESNCTAQEAQDDHHHCHPNTAGEEHIGFCRATVIRSHASLEKPKDEQFESDCVIFDVKKEHAHCYLIEQHVGKDSTSPGDNTNVLNLAHSHNHTRASHESHSDECVVGCPVAFVKKEVPTDISDHDTPSVKGCPGRVLHFNL*

>Vber_FETUA2b

MNSLVALLLLGQVIGSTLSSQVRGDIDCNDKDAKHWADIAVRYINEHKLHGYKQDLNVIKNIHILTWDRDLVAGYLKLNLLETECHVLDPTPVEKCTVRPQHNHTVEMDCDVKIIYNVMTLKDEVSVKCNSTTDSVENVQRNCPKCPILLPRNDPHVVDAVEYVLNKHNEKLSGHVYEVLEISRGQHTFEPEGFYVEFAIVQSNCTAQEAQDDHHHCHPNTAGEDHIGFCRATVFRSHASLEKPKDEQFESDCVMFDVKEGHTHSYLIEQHVGKDSTSPGDNTTVLNLAHSHNHTRASHEFHSDECVVECPVAFVKKEVPTDISDHHTPPAKGCPGRVLHFKL*

>Vlat_FETUA1

MNSLVALLLLGQIIGCTFSHHLPSHGDCNGEDAKKWAHLAVHYINKHTLHGYKQDLNIIKDIHVLPRRPHGKIVFLELELLETVCHVLDPTPVENCTVRPQHYHAVEGDCDVKIIHDEDVDNVVAAKCHSSPDSVEDVQQNCPTCPILLPLNDPHVVDAVEYVLNKHNEKLSGHVYEVLEISRGQHTFEPEGFYVEFAIVESNCTAQEAQNDHHHCHPNTAGEDHIGFCRANVIRSHASLEKPKDEQFESDCVIFDVKEGHSYSHLIEHHVGKYSTSPGHNNTVLNLAHSHNHTSASHESHSHEHVVEVPVAVAKREIPTNIPPHHTHPVNLCPGKVHHFKV*

>Vlat_FETUA2a

MNSLVALLLLSQIIGSTLSSQVRGDIDCNDKDAKDWAGKAVRYINEHKLHGYKQALNVIKNIHILPWDGDLVAIYLKLNFLETECHMLDPTPVENCTVRPQHNHAVKIDCDVKIIYDIETHKDDVFVKCNSTPDSVENVQQNCPKCPILLPQNDPHVVDAVEYVLNKHNEKLSGHVYEVLEISRGQHTVFSFCIRMAVMMVILGFLSSAVTLHNSKLHIETFRLKHIGFCRATVFRSHASLEKPKDEQYESDCVIFDVKGHAHCYLIEQHVGKDSTSPGDNTNVLNLAHSHNHTRASHESHSDECVVGCPVAFVKKEVPTDISDHDTPSVKGCPGRVLHFNL*

>Vlat_FETUA2b

MNSLVALLLLGQVIGSTLSSQVRGDIDCNDKDAKHWADIAVRYINEHKLHGYKQDLNVIKNIHILTWARDLVAGYLKLNLLETECHVLDPTPVEKCTVRPQHNHTVEMDCDVKIIYNVMTLKDEVFVKCNSTTDSVENVQRNCPKCPILLPRNDPHVVDAVEYVLNKHNEKLSGHVYEVLEISRGQHTFEPEGFYVEFAIVQSNCTAQEAQDDHHHCHPNTAGEDHIGFCRATVFRSHASLEKPKDEQFESDCVMFDVKEGHTHSYLIEQHIGKDSTSPGDNTTVLNLAHSHNHTRASHEFHSDECVVECPVAFVKKEVPTDISDHHTPPAKGCPGRVLHFKL*

>Afea_FETUA1

MNSLVALVLLAQIIGSTLSHHLPSHVDCNGEDAEKWADMAVHYINEHNVHGYKQALNVIKEIRVLPRRPRGEIVYLELELLETVCHVLDPTPVENCTVRPQHYHAVEGDCDVKIIHDEGVDKVVGAKCHSNPDSLEDVRQNCLKCPILLPLSDPHVVDSVEYVLNKHNEKLSDHVYEVLEISRGQHKYEPEAFYVEFAIVETNCTAQESHDDHHHCHPNTAGEAHIGFCRATVFRSHASLEKPKDEKIESDCVIFDVKEGHAHSHLIEHHVGKYSVSPGHNSIVLNLAHSHNHSSASHESHSHEHVTEVPVAVAKREVPTDIPHHHTHPVKGCPGKVHHFKL*

>Afea_FETUA2

MNSLVALVLLGQIIGSTLSSQVRGDLECDDKEAKEWAADAVRYINEHKLRGYKQALNVIKNILVVPWNGDLVAIFLKLNLLETECHVLDPTPVENCTVRPQHNHTVEMDCDAKIIFDIVTFKQDVFVKCNSTPDSLENVRRNCPKCPILLPLNDPHVVDSAEYVLNKHNEKLSDHVYEVLEISRGQHKYEPEAFYVEFAIVETNCTAQESHDDHHHCHPNTAGEDHIAFCRATVFRSHASLEKPKDEKFESDCVIFDVKEGHAHSHLIEQHESHSDECVAEALFAFVKKERPTDISDRHTHPVKGCPGKVHHFEL*

>Afea_FETUA3

MNSLVALVLLGHIIGSTLSFQLPGDMDCNTKGTKDWADIGVRYINEHKLHGYKQALNVIKIFRLLPSDGRSVIFYFKLNLLETECHVLDPTPVEKCTVRPQHNHAVEMDCNVKIIYDIVTFKDEVFVKCNSTPDSLENVRRNCPKCPILLPLSDPHVVDSAEYVLNKHNEKLSDHVYEVLEISRGQHKYEPEAFYVEFAIVETNCTAQESQDDHHHCHPNTAGEDHIGFCRATVFRSHASLEKPKDEKVESDCVIFDVKEGHAHSYLIEQHVGKYSVSSGHNSTVLNLAHSHNHTRPSHESHSDECVMECPVAFVKKEVPTDISDHDTPPVEGCPGRVLHFKL*

>Balt_FETUA1

MNSLVALVLLGQIIGSTLSHHLQSHVDCNGKDAEKWADMAVHYINEHNQHGYKCALNVINEIRLLPRRPHGTIVFLELKVLETVCHVLDPTPIENCTVRPQHYHAVGGDCDVKIIHEEGGDKVIGAKCYSDPDSVEDVRRNCPKCPILLNLNDPQVVDSVEYVLNKHNEKVSGHVYEVLEISRGQHKYEPEAYYVEFAIVEVNCTAQELHDDHHQCHPNTAGENHIGFCRGTVFRSHASLEKPKDEQFESDCVIFDVKDGHAHSHLIEHHVGKYSTSPGHNNTVLNLVHSHNHTSASHESHSHEHVAEVPVAVAKREVPKDIPHDHTHPVKLCPGKVHHFEL*

>Balt_FETUA2p

MNSLVALVLLGQIIGSTLSSQVRGDLECDEKDAKEWTDIGVRYINEHKLHGYKYALNVIKNIVVVPWDGDWVAVFLKLNLLETECHVLDPTPVKNCTVRTQHNHAVEMDCDVKIMFNVDTFKEDVFAKCHSTPDSVEDVRRNCPKCPILLPPNNPQVVDSVEYVLNKHNEKLSDHVYEVLEISRGQHKYEPEAYYVEFAIVEVNCTAQELHDDHHQCHPNTAGEDHIGLCRATVFRSHASLEKPKDEQFESDCVILHVKEGHAHSHLIQQHVEKESISPEHSNTAVNLVHPHNDTNTSHESHEHLAEVPVAFVKKELPKDISDRHTTPVKGCPGKVLHFQL*

>Balt_FETUA2q

MNSLVALVLLGQIIGSTLSSQVRGDLPCDDKDSKWWADIGVRYINEHKLHGYKYALNVIKNIVVTSRDRDRVAVFLKLNLLETECHVLDPTPVKNCAVRTQHNHAVEMDCDVNIMFNIETFKKDVFVKCHSTPDSVENVRRNCTKCPILLPPNNPQVVDSVEYVLNKHNEKVSGHVYEVLEISRGQHKYEPEAYYVEFAIVEVNCTAQEAHDDHHHCHPISAGEDHIAFCRATVFRSHASLEKPKDEQVESDCVILHVKKGHAHSHLIQQHVEKDSISPEHNNTALNLVHPHNDTSTSHESHEHLAEVPVAFVKKELSKDISDRHTTPVKGCPGKVLHFQL*

>Balt_FETUA2r

MNSLVALVLLGQIIGSTLSSQVRGDLQCDDEDAKEWTDRGVRYINEHKLHGYKYALNVIKNIVVVPLDGDWVAVFLKLNLLETECHVLDPNPFDNCTVRPQHNHAVEMDCDVKIMFNVDTFKEDVFAKCHSTPDSVENMQRNCPKCPILLPSNDPQVVDSAKYLLSKDNEKLFDQVYLYELLDISRGQHKYEPEAYYVEFAIVEVNCTAHELHDDHHQCHPNTAGEDHVALCIGTVFRSNASLEKLKDEKFEVDCFTILAKKGHAYSHLIQQLFEKKIISPEHNNTALNLIHPHNDTSTSHESHEHVAEVPVAFVKKELPKDISDHHTTPVKGCPGIVLHFEL*

>Balt_FETUA2s

MNSLVALVLLGQIIGSTLSSQVRGDLPCDDKDSKWWADIGVRYINEHKLHGYKYALNVIKNIVVTSRDHDRVAVFLKLNLLETECHVLDPTPVKNCAVRTQHNHAVEMDCDVNIMFNIETYKKDVFVKCHSTPDSVEDVRRNCTKCPILLPSNNPQVVDSVEYVLNKHNEQLSGHVYELLDISRGQHKYEPEAYYVEFAIVEVNCTAQEAHDDHHHCHPISAGEDHIAFCRATVFRSHASLEKPKDEQVESDCVILHVKKGHAHSHLIQQHVEKDSISPEHNNTALNLVHPHNDTSTSHESHEHLAEVPVAFVKKELSKDRSDRHTTPVKGCPGKVLHFQL*

>Bcot_FETUA1

MNSLVALVLLGQIIGSTLSHHLQSHVDCNGEDAEKWADMAVHYINEHNQHGYKCALNVINEIRLLPRRPHGTIVFLELKVLETVCHVLDPTPIENCTVRPQHYHAVGGDCDVKIIHEEGGDKVIGAKCYSDPDSVEDVRRNCPKCPILLSLNDPQVVDSVEYVLNKHNEKVSGHVYEVLEISRGQHKYEPEAYYVEFAIVEVNCTAQEAHDDHHQCHPNTAGENHIGFCRGTVFRSHASLEKPKDEQFESDCVIFDVKDGHAHSHLIEHHVGKYSTSPGHNNTVLNLVHSHNHTSASHESHSHEHVAEVPVAVAKREVPKDIPHDHTHPVKLCPGKVHHFEL*

>Bcot_FETUA2w

MNSLVALVLLGQIIGSTLSSQVRGDLECDEKDAKEWTDVGVRYINEHKLHGYKYALNVIKNIVVVPWDGDWVAVFLKLNLLETECHVLDPTPVKNCTVRTQHNHAVEMDCDVKIMFNVDTFKEDVFAKCHSTPDSVENVRRNCPKCPILLPSNNPQVVDSVEYVLNKHNEKLSGHVYEVLEISRGQHKYEPEAYYVEFAIVEVNCTAQEAHNDHHQCHPNTAGENHIAFCRATVFRSHASLEKPKDEQFESDCVILHVKQGHAHSHLIQQHVEKDSISPEHNNTALNLIHPHNDTSTSHESHEHVAEVPVAFVKKELPKDISDRHTTPVKGCPGKVLHFGL*

>Bcot_FETUA2x

MNSLVALVLLGQIIGSTLSSQVREDLECNDETAKWWTDIGVRYINEHKLHGYKYTLNVIKNIVVTSRDRDRVAVFLKLNLLETECHVLDPTPVKNCAVRTQHNHAVEMDCDVNIMFNVATFKEDVFAKCHSTPDSVENVRRNCTKCPILLPSNNPQVVDSVEYVLNKHNEQISGHVYEVLEISRGQHKYEPEAYYVEFAIVEVNCTAQEAHDDHHHCHPISAGEDHIAFCRATVFRSHASLEKPKDEQVESDCVILHVKKGHAHSHLIQQHVEKDSISPEHNNTALNLVHPHNDTSTSHESHEHLAEVPVAFVKKELSKDISDRHTTPVKGCPGKVLHFQL*

>Bcot_FETUA2y

MNFLVALVLLGQIIGSTLSSQVMGDLSCNDEDSRWWADVGVRYINEHKLHGYKYALSVIKNIIVVPWDGDWVAVFLKLNLLETKCHVLDPTPVKNCNVRTQHNHAVEMDCDVKIMFNVATFKEDVFAKCHSTPDSVEDVRRNCPKCSILLPSNHPQVVDSAEYVLNKHNEKLSDHVYEVLEISRGQHKYELEAYYVEFAIVEVNCTAQEAHDDHHQCHPNTAGENHIGFCRATVFRSHASLEKPKDEQFESDCVILHVKGHAHSHLIQQHVEKNSISAEHNNTALNFVHPHNDTRASHESHEHLAKVPVAFVKKELPKDISDRHTTPVFGCPGLRAVIQPY

>Bcot_FETUA2z

MNSLVALVLLGQIIGSTLSTQVRGDLQCDDKATKEWTDIGVRYINEHKLHGYKYALNVIKNIVVTSRDRDLVAVFLKLNLLETECHVLDPTPVKNCTVRTQHNHAVEMDCDVNIMFNVDTFKEDVFVKCHSTPDSVENVRRNCPKCPILLPSNNPQVVDSVEYVLNKHNEQLSDHVYEVLEISRGQHKYEPEAYYVEFAIVEVNCTAQEAHDDHHHCHPNTAGEDHIGFCRATVFRSHATLEKPKDEQFESDCVILHVKKGHAHSHLIQQHVEKESISPEHNNTALNLAHPHNDTSTSHESHEHLAKVPVAFVKKELSKDISDRHTTPVKGCPGKVHHFEL*

>Bcot_FETUA2za

MNSLVALMLLGQIIGSTLSSQVRGDLECNDKDAKEWTDVGVRYINEHKLHGYKYALNVIKNIVVTSRDRDWVAVFLKLNLLETECHVLDPTPVKNCAVRTQHNHAVEMDCDVNIMFNVDTFKEDVFVKCHSTPDSVENVRRNCPKCPILLPPNHPQVVDSVEYVLNKHNEKLSGHVYEVLEISRGQHKYEPEAYYVEFAIVEVNCTAQEAHDDHHHCHPNTAGEDHIGFCRATVFRSHATLEKPKDEQFESDCVILHVKKGHAHSHLIQQHVEKESISPEHNNTALNLAHPHNDTSTSHESHEHLAKVPVAFVKKELSKDISDRHTTPVKGCPGKVHHFEL*

>Bfon_FETUA1

MNSLVALVLLGQIIGSTLSHHLQSHVDCNGEDAEKWADMAVHYINEHNQHGYKCALNVINEIRLLPRRPHGTIVFLELKVLETVCHVLDPTPIENCTVRPQHYHAVGGDCDVKIIHEEGGDKVIGAKCYSDPDSVEDVRRNCPKCPILLSLNDPQVVDSVEYVLNKHNEKVSGHVYEVLEISRGQHKYEPEAYYVEFAIVEVNCTAQEAHDDHHQCHPNTAGENHIGFCRGTVFRSHASLEKPKDEQFESDCVIFDVKDGHAHSHLIEHHVGKYSTSPGHNNTVLNLVHSHNHTSASHESHSHEHVAEVPVAVAKREVPKDIPHDHTHPVKLCPGKVHHFEL*

>Bfon_FETUA2g

MNSLVALVLLGQIIGSTLSSQVRGDLECDEKDAKEWTDVGVRYINEHKLHGYKYALNVIKNIVVVPWDGDWVAVFLKLNLLETECHVLDPTPVKNCTVRTQHNHAVEMDCDVKIMFNVDTFKEDVFAKCHSTPDSVENVRRNCPKCPILLPSNNPQVVDSVEYVLNKHNEKLSGHVYEVLEISRGQHKYEPEAYYVEFAIVEVNCTAQEAHNDHHQCHPNTAGENHIAFCRATVFRSHASLEKPKDEQFESDCVILHVKQGHAHSHLIQQHVEKDSISPEHNNTALNLIHPHNDTSTSHESHEHVAEVPVAFVKKELPKDISDRHTTPVKGCPGKVLHFGL*

>Bfon_FETUA2h

MNSLVALVLLGQIIGSTLSSQVREDLECNDETAKWWTDIGVRYINEHKLHGYKYTLNVIKNIVVTSRDRDWVAVFLKLNLLETECHVLDPTPVKNCAVRTQHNHAVEMDCDVNIMFNIETYKKDVFVKCHSTPDSVENVRRNCTKCPILLPSNNPQVVDSVEYVLNKHNEQISGHIYEVLEISRGQHKYEPEAYYVEFAIVEVNCTAQEAHDDHHQCHPNTAGENHIAFCRATIFRSHASLEKPKDEQVESDCVILHVKKGHAHSHLIQQHVEKDSISPEHNNTALNLVHPHNDTSTSHESHEHLAEVPVAFVKKELSKDISDRHTTPVKGCPGKVLHFQL*

>Bfon_FETUA2i

MNSLVVLVLLGQIIGSTLSSQVRGDLECDEKDAKEWTDVGVRYINEHKLHGYKYALNVIKNIVVVPWDGDWVAVFLKLNLLETECHVLDPTPVKNCTVRTQHNHAVEMDCDVKIMFNVDTFKEDVFAKCHSTPDSVENVRRNCPKCPILLPSNHPQVVDSVEYVLNKHNEKLSGHVYEVLEISRGQHKYEPEAYYVEFAIVEVNCTAQEAHDDHHQCHPNTAGEDHIGFCRATVFRSHASLEKPKDEQFESDCVILHVKEGHAHSHLIQQHVEKDSISPGHNNTALNLVHPHNYTGTSTSHESHEHLAEVPVAFVKKELPKDISDRHTTPVKGCPGKVHHFEL*

>Bfon_FETUA2j

MNSLVALVLLGQIIGSTLSTQVRGDLQCDDKATKEWTDIGVRYINEHKLHGYKYALNVIKNIVVTSRDRDLVAVFLRLNLLETECHVLDPTPVKNCTVRTQHNHAVEMDCDVNIMFNVDTFKEDVFVKCHSTPDSVEDVRRNCPKCPILLSLNDPQVVDSVEYVLNKHNEKLSGHVYEVLEISRGQHKYEPEAYYVEFAIVEVNCTAQEAHDDHHHCHPNTAGEDHIGFCRATVFRSHASLEKPKDEQFESDCVILHVKKGHAHSHLIQQHVEKESISPEHNNTALNLAHPHNDTSTSHESHEHLAKVPVAFVKKELSKDISDRHTTPVKGCPGKVLHFGL*
